# Targeted siRNA screening coupled with unbiased proteomics identify potassium signaling as a critical regulator of pigmentation

**DOI:** 10.64898/2026.11.13.766383

**Authors:** Kriti Ahuja, Ishita Panthri, Abhishek Tanwar, Anuradha Jadon, Ankit Biswas, Tushar Kanti Maiti, Rajender K. Motiani

## Abstract

Ion channels are emerging as critical regulators of cellular signaling, influencing diverse pathophysiological processes and offering promising opportunities for therapeutic intervention. However, the contribution of ionic homeostasis to human skin pigmentation remains poorly understood. Here, we identify potassium (K^+^) dynamics as a critical regulator of pigmentation and uncover K^+^ channel Kir7.1 as a molecular link between K^+^ and cyclic adenosine monophosphate (cAMP) signaling. A targeted siRNA screen identified six previously unrecognized regulators of pigmentation, which we validated using mouse cell line systems, primary human melanocytes, and an *in vivo* zebrafish model. Unbiased proteomic analyses of two prominent screening hits, Rhot1 and VDAC3, led us to identify Kir7.1, a plasma membrane-localized K^+^ channel, as a key determinant of pigmentation. Integrating cellular, molecular, ultrastructural, and microscopy approaches, we demonstrate functional crosstalk between the hits and Kir7.1 channel activity. Mechanistically, Kir7.1 promotes the canonical cAMP–protein kinase A (PKA) pathway, establishing this channel as a molecular node that couples K^+^ dynamics to the classical melanogenic pathway. Notably, Kir7.1 is differentially expressed in skin pigmentary disorders such as vitiligo and melasma, highlighting its therapeutic potential. Taken together, our findings establish Kir7.1 as an important regulator of skin pigmentation and reveal an unanticipated connection between K^+^ homeostasis, mitochondrial Ca^2+^ handling toolkit, and cAMP-dependent melanogenesis. This work provides a foundation for exploring K^+^ signaling as a potential therapeutic avenue for human pigmentary disorders.

**Graphical Abstract:** 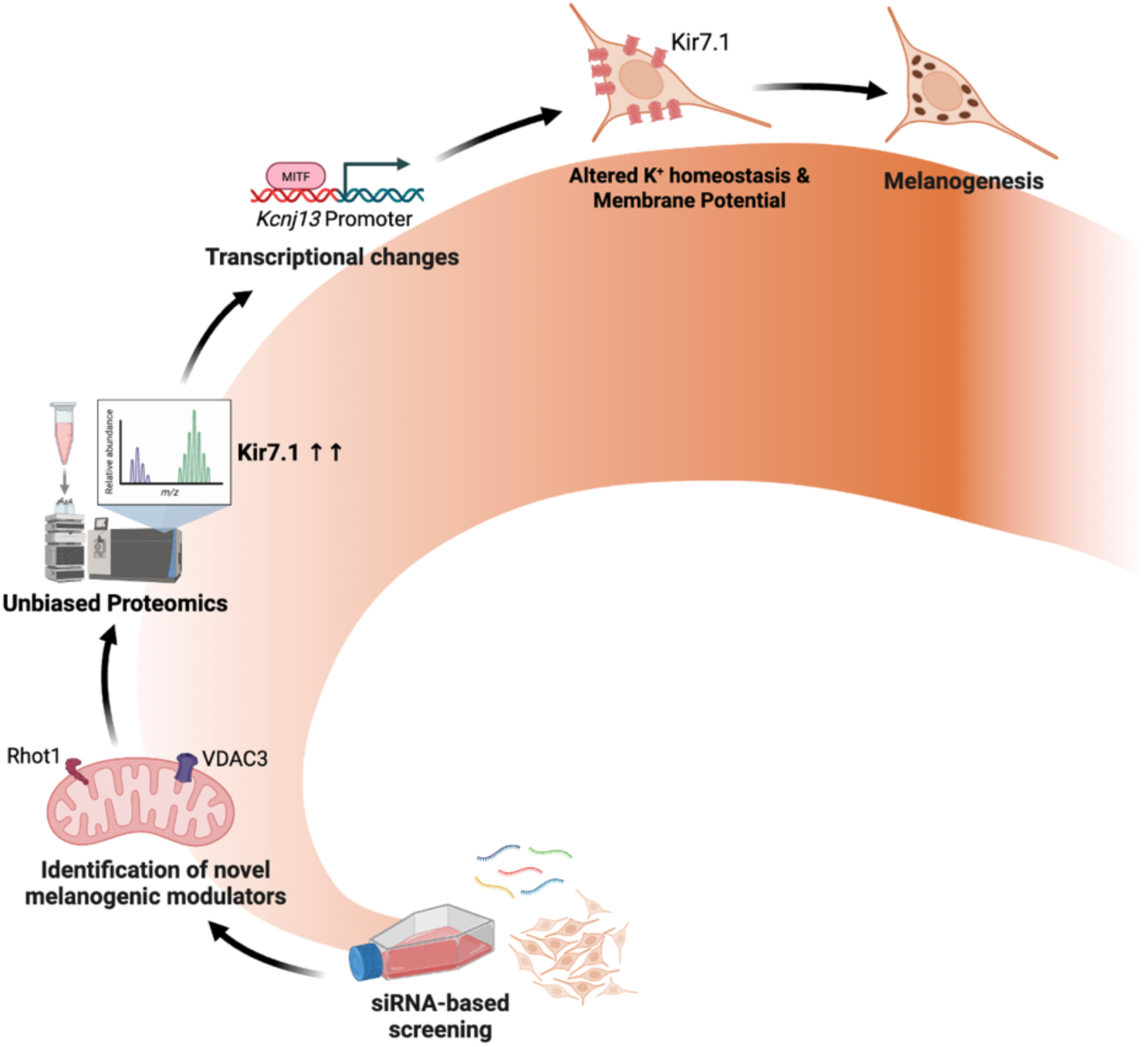

## Introduction

Ion homeostasis is one of the most fundamental determinants of cellular function. Cells maintain large ion concentration gradients across extracellular milieu and cytosol as well as endomembrane organelles and cytosol [1]. This facilitates rapid information exchange via controlled ion flux in response to cellular signals. By regulating selective flow of ions across lipid bilayers, they control electrical excitability, muscle contraction, secretion, volume regulation, govern gene expression, proliferation and cell death [2]. Their dysfunction results in pathologies such as cardiomyopathy, cancer, epilepsy, sensory and metabolic disorders [2]. Therefore, in recent years, ion channels have evolved as attractive targets for drug discovery. Ionic exchange across the lipid bilayers is mediated by ion channels and transporters distributed across the endomembrane system, including mitochondria, endoplasmic reticulum (ER), Golgi apparatus, and lysosomes [3]. Precise regulation of ion homeostasis and dynamics across these organelles is indispensable for cellular function.

Mitochondria are multifunctional organelles that serve as central regulators of cellular homeostasis [4]. Their functions extend beyond their classical role as the ‘powerhouse’ of the cell. Mitochondria play a critical role in calcium (Ca²⁺) homeostasis, redox balance, apoptosis, autophagy, cell bioenergetics, intracellular signaling and metabolic adaptations [5]. Disruption of mitochondrial homeostasis can impair cellular function, induce metabolic imbalance, trigger oxidative stress, activate the stress response, and ultimately trigger cell death pathways. Therefore, mitochondrial dysfunction can contribute to the pathogenesis of various disorders [3,6]. Mitochondrial Ca²⁺ homeostasis is one of the most crucial processes regulated by mitochondria, serving as a central determinant of cellular physiology by integrating bioenergetics, metabolism and signaling processes [5–8]. Perturbations in mitochondrial Ca^2+^ dynamics are implicated in numerous disorders, including cancer, neurodegenerative diseases, cardiovascular diseases, metabolic syndromes, and melanogenic disorders [6].

The transfer of Ca^2+^ into mitochondria represents a finely orchestrated inter-organelle process. Ca^2+^ released from intracellular reservoir organelles is funneled across the double membrane system of mitochondria i.e. the outer mitochondrial membrane (OMM) and the inner mitochondrial membrane (IMM) [4]. Voltage-dependent anion channels (VDACs) on the OMM represent the primary molecular interface that ensures high bidirectional Ca^2+^ permeability across the OMM, allowing rapid equilibration of cytosolic and intermembrane space (IMS) Ca^2+^ levels [4,9,10]. The three isoforms of VDACs (VDAC1, VDAC2 and VDAC3) share about ∼70% sequence similarity in mammals, with VDAC1 and VDAC2 being the most abundant isoforms and VDAC3 being the least abundant [9,11]. Literature highlights the role of VDACs in regulating metabolism, ion transport, apoptosis and cell death pathways [9,11–13]. The OMM also harbors mitochondrial Rho GTPases 1 and 2 (Rhot1 and Rhot2), also known as MIRO1 and MIRO2, which regulate mitochondrial Ca^2+^ dynamics, mitochondrial transport and spatial positioning, ER-mitochondria communication, and mitophagy [14,15]. On the IMM, Ca^2+^ influx from IMS to the matrix is predominantly mediated by the MCU macromolecular complex, formed by subunits-MCU (the pore), MCUb (dominant-negative subunit), MICU1, 2 and 3 (gatekeepers), EMRE (scaffold protein) and MCUR1 (regulatory subunit) [10,16,17]. Recent evidence indicates the existence of MCU-independent mitochondrial Ca^2+^ uptake mechanisms, as MCU-deficient mitochondria still exhibit some Ca^2+^ storage [10]. MCU knockout mice exhibit unaltered basal cardiac parameters, suggesting compensatory mechanisms that enable their survival in the absence of MCU [17]. Some of the MCU-independent mitochondrial Ca^2+^ uptake mechanisms reported involve mitochondrial uncoupling proteins 2 and 3 (UCP2 and UCP3) [18]. To maintain mitochondrial Ca^2+^ homeostasis, regulated Ca^2+^ extrusion from the matrix is equally essential. Mitochondria can extrude Ca^2+^ from the matrix via Na^+^/Ca^2+^ exchangers (mNCX), the mitochondrial H^+^/Ca^2+^ exchangers (mHCX) and the non-specific mitochondrial permeability transition pore complex (mPTP) [4,10]. The IMM-localized Leucine zipper EF-hand-containing transmembrane protein 1 (LETM1) channels also mediate mitochondrial Ca^2+^ dynamics, but their mechanism remains unresolved and controversial [4,19]. This coordinated network of multiple channels and transporters underscores the sophistication of mitochondrial Ca²⁺ signaling mechanisms.

One of the key processes regulated by cellular Ca^2+^ signaling is skin pigmentation [20–23]. Melanogenesis, the process of synthesis of the skin pigment melanin, is a multifactorial physiological phenomenon that imparts photoprotection to the skin, thereby preventing UV-induced DNA damage, mutagenesis, and the development of skin cancer and pigmentary anomalies [20]. The melanin is synthesized in specialized neuro-ectoderm derived dendritic cells called melanocytes [20]. Melanocytes synthesize two forms of melanin-red-yellow pheomelanin and brown-black eumelanin, and their relative ratio determines the diverse pigmentation phenotypes observed in human skin. Melanogenesis begins with tyrosinase, the rate-limiting enzyme that converts tyrosine to DOPA and dopaquinone, with dopachrome tautomerase (DCT/TYRP2) then driving eumelanin synthesis. Melanin is synthesized within lysosome-related organelles called melanosomes where PMEL17 or GP100 is involved in scaffolded fibril assembly, enzyme loading, and pigment accumulation. Finally, mature melanosomes are transferred to keratinocytes to produce visible skin pigmentation [20,24] .

Recently, mitochondria and the associated signaling cascades have emerged as key regulators of melanogenesis. We have previously shown that mitochondria-localized proteins Mitofusin2 (Mfn2) and Mitochondrial Calcium Uniporter (MCU) are negative and positive regulators of pigmentation, respectively [16,25]. In the latter study, we show that the MCU-NFAT2-Keratin5-mediated feedback loop orchestrates pigmentation by regulating mitochondrial Ca^2+^ homeostasis [16]. Therefore, mitochondrial Ca^2+^ signaling machinery is crucial for ensuring optimal melanogenesis. However, the mitochondrial Ca^2+^ toolkit comprises numerous Ca^2+^-sensitive proteins and channels that ensure efficient ion homeostasis, but their functional significance in pigmentation biology remains elusive.

In this study, we utilized an unbiased siRNA-mediated screening approach to identify mitochondria-localized Ca^2+^ signaling proteins that potentially regulate pigmentation. Using multiple model systems namely mouse B16 low density (LD) model, αMSH (αMelanocyte Stimulating Hormone)-induced physiological melanogenesis, primary human melanocytes and zebrafish *in vivo* models-we reveal that mitochondrial Ca^2+^ regulator proteins Rhot1, VDAC1, VDAC2 and VDAC3 negatively modulate pigmentation. Mechanistically, unbiased proteomics analyses of Rhot1 and VDAC3 silenced cells identified a cell membrane localized K^+^ efflux channel, Kir7.1 as a key downstream pigmentation modulator. Further, we identify K^+^ homeostasis as a novel mechanism governing melanogenesis. We also show that Kir7.1 expression is regulated downstream of MITF transcription factor suggesting that Kir7.1 could be an essential player contributing to the classical αMSH-mediated pigmentation cascade. Finally, we show that Kir7.1 is differentially regulated in hypopigmentary and hyperpigmentary skin disorders, highlighting its potential as a disease biomarker and/or therapeutic target for managing pigmentary disorders.

## Results

### Unbiased siRNA screens targeting mitochondrial Ca^2+^ handling machinery identify novel modulators of pigmentation

The functions of mitochondrial Ca^2+^ handling machinery and dynamics in pigmentation remain poorly appreciated. Recently, we revealed that Mitochondrial Ca^2+^ Uniporter (MCU) complex mediated mitochondrial Ca^2+^ uptake can augment melanogenesis via transcriptional regulation of keratins [16]. To comprehensively dissect the role of mitochondrial Ca^2+^ handling proteins in mediating pigmentary responses, we performed unbiased siRNA-based screens targeting these proteins using a well-established B16 mouse melanoma cell line based low-density (LD) pigmentation model [16,23,26,27]. LD pigmentation model utilizes non-pigmented B16 cells seeded at a very low cell density of 100 cells/cm^2^. This results in cell-autonomous and synchronous pigmentation induction over a time course of 6-7 days as depicted in the schematic diagram (**Fig. 1A**). We quantitated the increase in pigmentation from LD day 0 to day 4 to day 7 in the LD pigmentation model by analyzing the mean pixel intensity of the pellets using FIJI software (**Fig. 1B**). The LD pigmentation model closely recapitulates the pigmentary response in primary human melanocytes [16,23,26,28]. We identified candidate proteins for screening using the mouse MitoCarta3.0 database, a library of nuclear and mtDNA genes with strong evidence of mitochondrial localization [29]. We employed the following selection criteria for target proteins: (i) presence of EF-hand domain(s) responsible for Ca^2+^ binding and/or (ii) they mediate Ca^2+^ transport across inner or outer mitochondrial membrane (**Fig. 1C**). From the list of compiled mitochondrial Ca^2+^ handlers, 7 proteins belong to the MCU complex (MCU, MCUb, MICU1, MICU2, MICU3, EMRE, and MCUR1), which has been previously screened in our lab and recently reported to regulate melanogenesis [16]. We then subjected the 17 other shortlisted candidate proteins to robust LD pigmentation model-based siRNA screening to identify if they modulate the pigmentation phenotype. We plated the B16 mouse melanoma cells at LD (Day 0), followed by siRNA transfections on Day 3 to induce targeted protein knockdown. The LD experiments were terminated on Day 6, and cell pellets were harvested to evaluate changes in pigmentation phenotypes (**Fig. 1D, S1A**).

**Figure 1.**
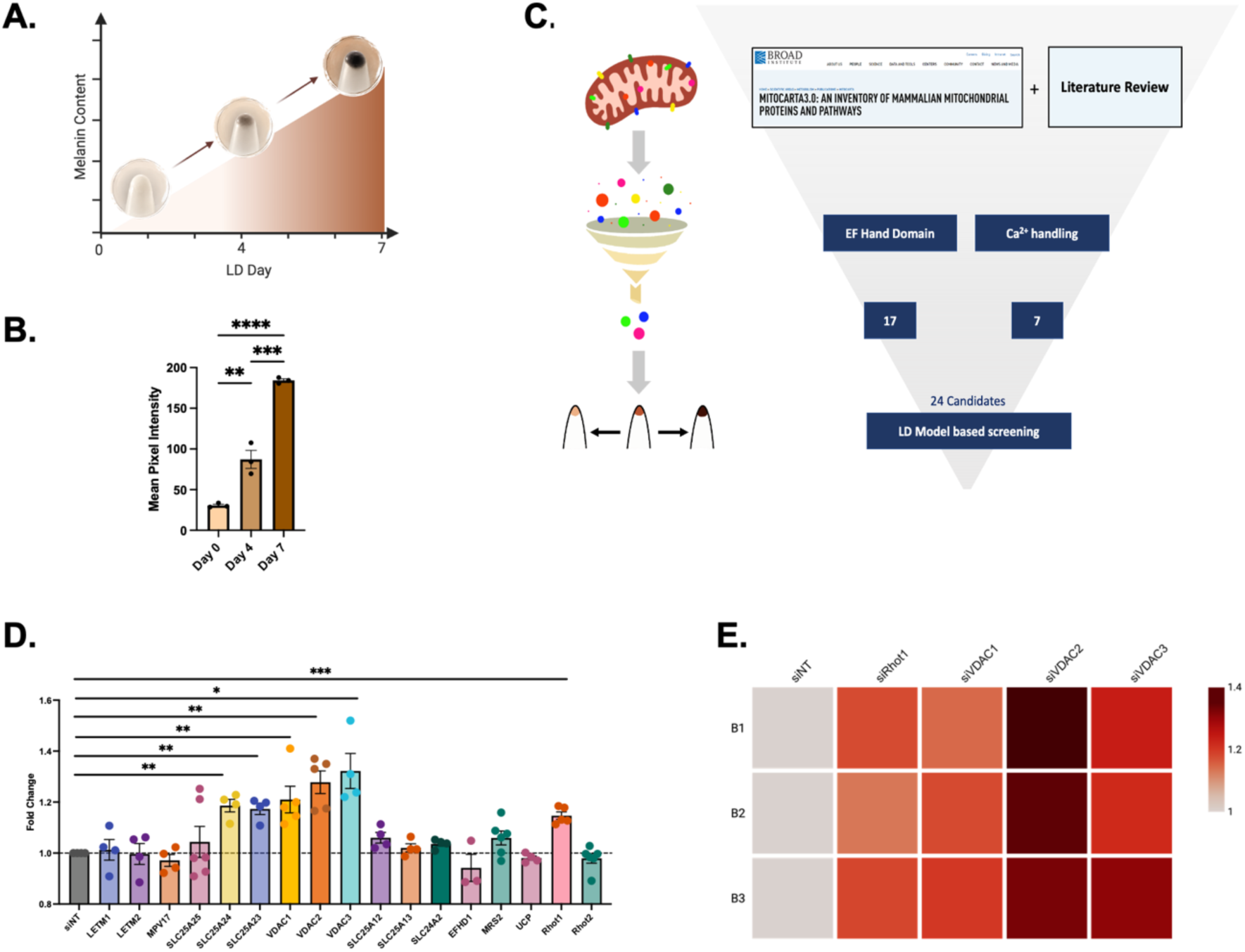
Unbiased siRNA screens targeting mitochondrial Ca^2+^ handling machinery identify novel modulators of pigmentation. A. B16 cell pellet pictures of the gradual increase in pigmentation from Day 0 to Day 4 and Day 7 in the B16 LD model system. B. Mean pixel intensity quantification of the pigmentation phenotype on LD Day 0, Day 4 and Day 7 using FIJI (N=3). C. Schematic representation of the workflow for selecting target candidates for LD model-based screening. D. FIJI-based pigmentation phenotype quantification of siRNA transfected B16 cells on LD day 6 as compared to siNT control (N=3-6). E. Heatmap of pigmentary phenotype changes in siRNA-transfected cells in LD model system (N= 3). Data presented are mean ± SEM. For statistical analysis, ordinary one-way ANOVA was performed for panel B and one sample t and Wilcoxon test was performed for panel D, using GraphPad Prism software. Here, *p<0.05; ** p<0.01; *** p<0.001; and **** p<0.0001. The data underlying panel D are presented in Fig. S1 Data.

For this study, we focused on two categories of hits obtained from our LD model-based screening i.e. Rhot1 (also known as Miro1) and Voltage Dependent Ion Channels (VDACs), silencing of which resulted in hyperpigmentation as compared to the control condition (**Fig. 1E**). These hits are individually discussed in detail in the subsequent sections.

### Rhot1 negatively regulates melanogenesis *in vitro*

To understand the role of Rhot1 in melanogenesis, we performed siRNA-based silencing of Rhot1 in B16 cells and observed a significant decrease in Rhot1 mRNA levels in siRhot1 cells compared with the non-targeting siRNA (siNT) control (**Fig. 2A**). Next, we investigated the role of Rhot1 in B16 LD melanogenesis. Silencing of Rhot1 on day 3 of LD model system using siRNA and termination on day 6 resulted in hyperpigmentation in siRhot1 cells compared to the siNT control as represented in the cell pellet images (**Fig. 2B**). We quantitated the increase in pigmentation upon Rhot1 knockdown by mean pixel intensity analysis of the cell pellets using FIJI software (**Fig. 2C**). We next examined the expression of melanosome structural protein GP100 or Pre-melanosome Protein 17 (PMEL17) and principal melanogenic enzymes – Dopachrome tautomerase (DCT) and tyrosinase upon Rhot1 silencing. Rhot1 knockdown resulted in substantial increase in the expression of GP100, DCT and tyrosinase (**Fig. 2D-2F and S2A-C**). We also assessed tyrosinase activity, a rate-limiting enzyme in the melanogenic cascade, using the Dopachrome generation (DOPA) assay, in which the enzymatic conversion of L-DOPA to dopachrome by tyrosinase results in the development of brownish-black bands on native gels. We observed that tyrosinase activity was also significantly enhanced in Rhot1 knockdown cells (**Fig. 2G, S2D**).

**Figure 2.**
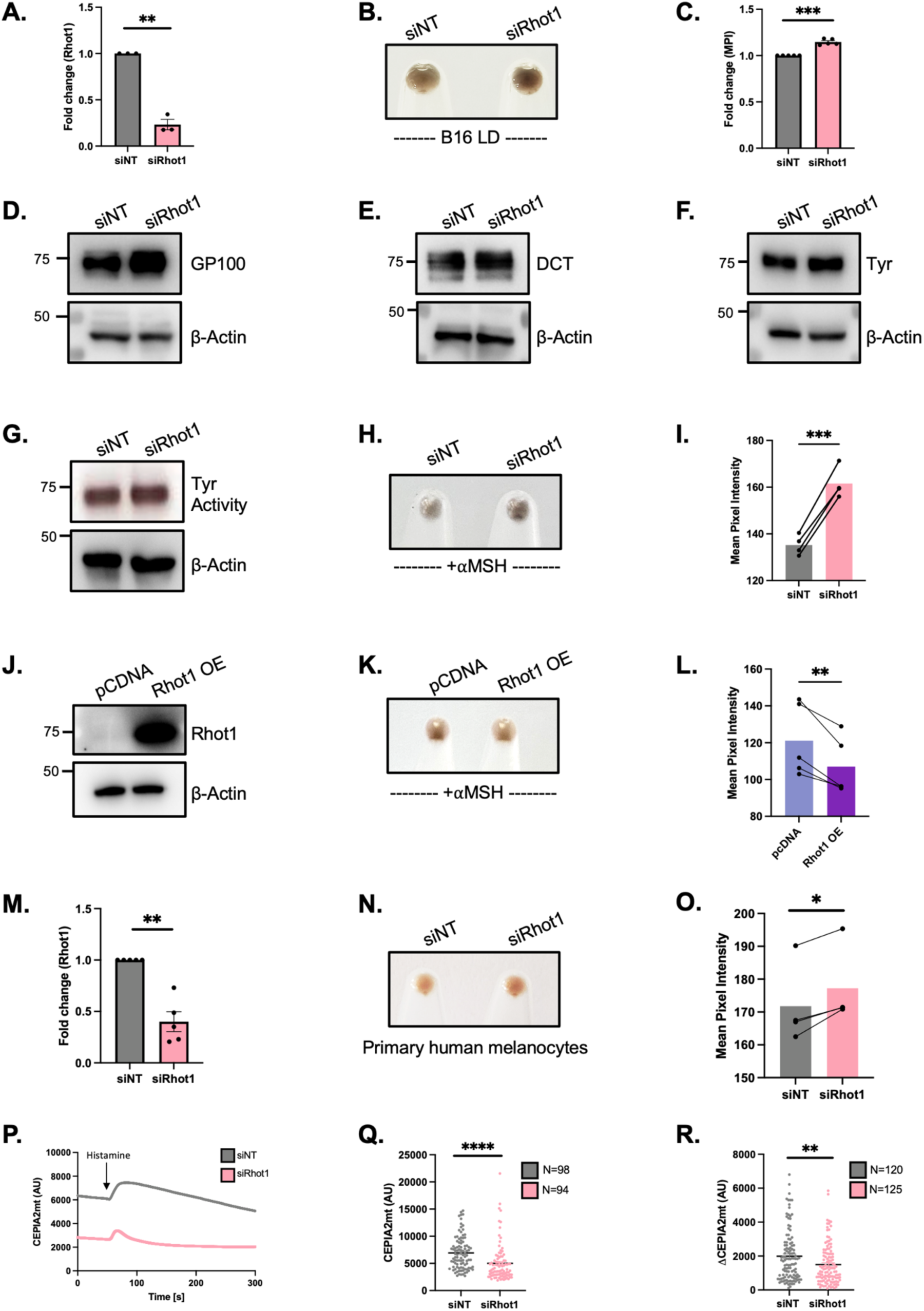
Rhot1 is a negative modulator of melanogenesis. A. qRT-PCR analysis confirming siRNA-induced silencing of Rhot1 on LD day 6 in B16 cells (N=3). B. Representative pellet pictures comparing LD day 6 pigmentation phenotype of siNT control and siRhot1. C. FIJI-based pigmentation phenotype quantification of siNT and siRhot1 B16 cells on LD day 6 (MPI=Mean Pixel Intensity, N=5). D. Representative western blot showing GP100 expression in siNT and siRhot1 LD day 6 cells (N=5). E. Representative western blot showing DCT expression in siNT and siRhot1 LD day 6 cells (N=3). F. Representative western blot showing tyrosinase expression in siNT and siRhot1 LD day 6 cells (N=3). G. Representative in-gel image of DOPA assay of siNT and siRhot1 LD day 6 cells (N=4). H. Representative pellet pictures comparing pigmentation phenotype of siNT control vs siRhot1 in HD-MSH pigmentation experiment. I. FIJI-based mean pixel intensity quantification of pigmentation phenotype in siNT and siRhot1 B16 cells in HD-MSH system (N=4). J. Representative western blot validating Rhot1 over-expression (OE) in Rhot1 OE condition compared to pcDNA3.1 control (N=3). K. Representative pellet pictures comparing pigmentation phenotype of control vs Rhot1 OE in HD-MSH B16 pigmentation system (N=5). L. FIJI-based mean pixel intensity quantification of pigmentation phenotype of control vs Rhot1 OE in HD-MSH B16 pigmentation system (N=5). M. qRT-PCR analysis confirming siRNA-based knock-down of Rhot1 in primary human melanocytes (N=5). N. Representative pellet pictures comparing pigmentation phenotype of siNT control vs siRhot1 in primary human melanocytes. O. FIJI-based mean pixel intensity quantification of pigmentation phenotype in siNT and siRhot1 B16 cells in primary human melanocytes (N=4). P. Representative mitochondrial Ca^2+^ imaging traces of CEPIA2mt in siNT control and siRhot1 B16 cells stimulated with 50µM histamine. Q. Quantitation of resting mitochondrial Ca^2+^ with CEPIA2mt in siNT control and siRhot1 B16 cells where “N” denotes the number of ROIs. R. Quantitation of mitochondrial Ca^2+^ uptake by calculating increase in CEPIA2mt signal (ΔCEPIA2mt) in siNT control and siRhot1 B16 cells stimulated with 50µM histamine where “N” denotes the number of ROIs. Data presented are mean ± SEM. For statistical analysis, one sample t and Wilcoxon test was performed for panels A, C and M, ratio paired t-test was performed for panel I, L and O and unpaired Student’s t-test was performed for panel Q and R, using GraphPad Prism software. Here, *p<0.05; ** p<0.01; *** p<0.001; and **** p<0.0001. The data underlying panels D, E, F, G, and J are presented in Fig. S2 Data.

Next, we substantiated the role of Rhot1 in physiological melanogenesis. We used αMSH to physiologically stimulate pigmentation in B16 cells plated at high density (HD) and silenced the gene of interest. Going forward, this model is referred as HD-MSH pigmentation system. In this model as well, we observed a robust increase in pigmentation upon siRNA-mediated knockdown of Rhot1 as compared to the siNT control (**Fig. 2H, 2I**). Next, we overexpressed Rhot1 in B16 cells using the pRK5-myc-Miro1 plasmid and validated Rhot1 overexpression by performing western blotting (**Fig. 2J, S2E**). We examined the effect of ectopic Rhot1 expression on αMSH stimulated pigmentation in B16 cells and observed a marked reduction in melanogenesis (**Fig 2K**, **2L**). We further corroborated the role of Rhot1 in regulating pigmentation in primary human melanocytes by transiently silencing Rhot1. We first characterized the Rhot1 siRNA and observed significant reduction (around 60%) in Rhot1 mRNA levels post transfection in human melanocytes (**Fig. 2M**). Consistent with the previous observations, Rhot1 knockdown in primary human melanocytes increased melanogenesis (**Fig. 2N, 2O**). Taken together, these loss of function and gain of function studies in B16 cells along with Rhot1 silencing in primary melanocytes demonstrate that Rhot1 is a critical modulator of melanogenesis.

Next, we investigated the impact of Rhot1 on mitochondrial Ca^2+^ homeostasis. We measured the basal mitochondrial Ca^2+^ levels as well as the mitochondrial Ca^2+^ uptake using cells transfected with mitochondria-specific genetically encoded Ca^2+^-measuring organelle-entrapped protein indicators (CEPIA2mt) [30]. Basal mitochondrial Ca^2+^ levels are indicated by the basal fluorescent signal of the genetic probe while histamine stimulation is used to study mitochondrial Ca^2+^ uptake. Histamine is a physiological agonist that mobilizes Ca^2+^ from the intracellular stores i.e. Endoplasmic Reticulum (ER) and stimulates the mitochondrial Ca^2+^ uptake [16] . Importantly, histamine exposure promotes melanogenesis by remodeling Ca^2+^ signaling and maintaining sustained Ca^2+^ influx in the cell [16,31,32]. Our live-cell mitochondrial Ca^2+^ imaging data in B16 cells reveal that Rhot1 silencing reduces both the basal mitochondrial Ca^2+^ levels and mitochondrial Ca^2+^ uptake (**Fig. 2P-2R**). Our previous study had demonstrated that mitochondrial Ca^2+^ levels are positively associated with melanogenesis [16]. Interestingly, we here observed an inverse relationship between the mitochondrial Ca^2+^ levels and the melanogenic outcome.

Collectively, results across three independent melanogenesis models: B16 LD model, αMSH induced pigmentation model and primary human melanocytes strongly suggest that Rhot1 is a negative modulator of pigmentation albeit being a positive regulator of mitochondrial Ca^2+^ levels.

### VDACs negatively regulate melanogenesis *in vitro*

Our robust LD model-based screening experiments implicated the role of VDAC1, VDAC2 and VDAC3 in melanogenesis (**Fig.1**). Consistent with our findings, two recent studies independently identified VDAC1 as a negative regulator of melanogenesis in [33,34]. These independent investigations validate the reliability and robustness of our siRNA screening strategy. However, VDAC2 and VDAC3 isoforms have not been previously implicated in melanogenesis. To delineate the functional significance of all VDAC isoforms in melanogenesis, we conducted siRNA-mediated knockdown experiments in B16 LD model system. First, we characterized the siRNAs targeting VDAC1, VDAC2, and VDAC3 and observed robust, isoform-specific reductions in VDAC1/2/3 mRNA levels in siRNA-transfected B16 LD-day 6 cells (**Fig. 3A-3C**). Next, we examined the phenotypic response upon silencing of VDAC1/2/3 isoforms in B16 LD model system. We observed a robust increase in melanogenic response in siVDAC1, siVDAC2, and siVDAC3 conditions compared with their respective siNT controls (**Fig. 3D-3F**). Mean pixel intensity-based phenotype quantifications of the siVDAC-induced hypermelanogenic response are shown beneath the pellet pictures (**Fig. 3G-3I**). We further examined the effects of VDAC1/2/3 knockdown on the expression of melanogenic protein DCT and on tyrosinase enzyme activity using DOPA assay. We observed that DCT protein expression was markedly increased upon silencing VDAC1, VDAC2 and VDAC3 (**Fig. 3J-3L and S3A-S3C**). This was also accompanied by increased tyrosinase activity upon knockdown of all three VDAC isoforms (**Fig. 3M-3O and S3D-S3F**). Our data clearly indicates that all three VDAC isoforms-VDAC1, VDAC2 and VDAC3 can specifically modulate the pigmentation phenotype, rather than just VDAC1 as reported earlier.

**Figure 3.**
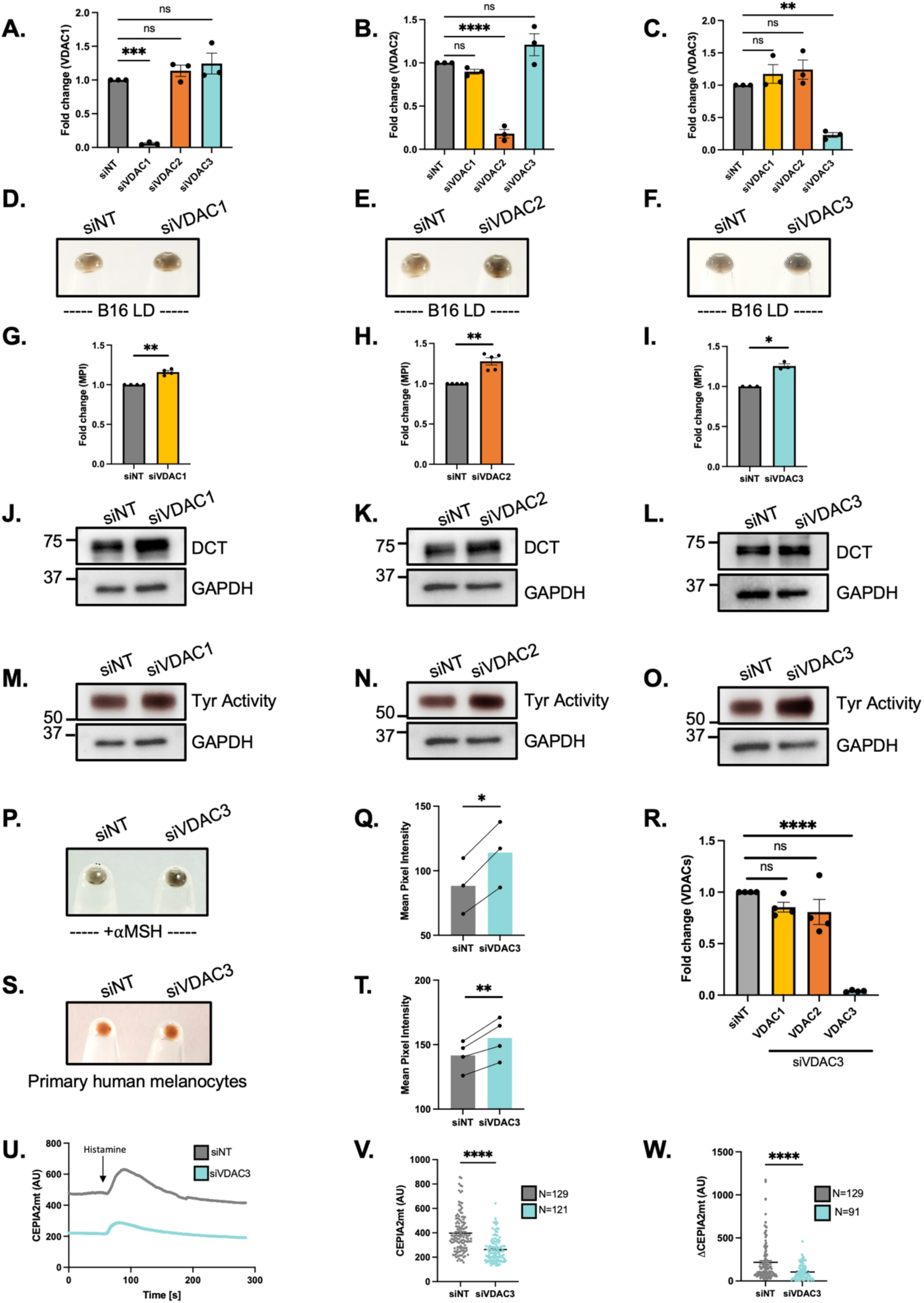
VDACs are negative modulators of melanogenesis. A. qRT-PCR analysis confirming siRNA-induced knock-down of VDAC1 in LD day 6 B16 cells, while VDAC2 and VDAC3 mRNA levels are unchanged (N=3). B. qRT-PCR analysis confirming siRNA-induced knock-down of VDAC2 in LD day 6 B16 cells, while VDAC1 and VDAC3 mRNA levels are unchanged (N=3). C. qRT-PCR analysis confirming siRNA-induced knock-down of VDAC3 in LD day 6 B16 cells, while VDAC1 and VDAC2 mRNA levels are unchanged (N=3). D. Representative pellet pictures comparing LD day 6 pigmentation phenotype of siNT control and siVDAC1 B16 cells. E. Representative pellet pictures comparing LD day 6 pigmentation phenotype of siNT control and siVDAC2 B16 cells. F. Representative pellet pictures comparing LD day 6 pigmentation phenotype of siNT control and siVDAC3 B16 cells. G. Mean pixel intensity (MPI) -based pigmentation phenotype quantification of siNT and siVDAC1 B16 cells on LD day 6 (N=4). H. Mean pixel intensity-based pigmentation phenotype quantification of siNT and siVDAC2 B16 cells on LD day 6 (N=5). I. Mean pixel intensity-based pigmentation phenotype quantification of siNT and siVDAC3 B16 cells on LD day 6 (N=3). J. Representative western blot showing expression of DCT in siNT control and siVDAC1 LD day 6 cells. K. Representative western blot showing expression of DCT in siNT control and siVDAC2 LD day 6 cells. L. Representative western blot showing expression of DCT in siNT control and siVDAC3 LD day 6 cells. M. Representative western blot showing tyrosinase activity in siNT control and siVDAC1 LD day 6 cells. N. Representative western blot showing tyrosinase activity in siNT control and siVDAC2 LD day 6 cells. O. Representative western blot showing tyrosinase activity in siNT control and siVDAC3 LD day 6 cells. P. Representative pellet pictures comparing pigmentation phenotype of siNT control vs siVDAC3 in HD-MSH pigmentation experiment. Q. FIJI-based mean pixel intensity quantification of pigmentation phenotype in siNT and siVDAC3 B16 cells in HD-MSH system (N=3). R. qRT-PCR analysis confirming siRNA-induced knock-down of VDAC3 in primary human melanocytes, while VDAC1 and VDAC2 mRNA levels are unchanged (N=4). S. Representative pellet pictures comparing pigmentation phenotype of siNT control vs siVDAC3 in primary human melanocytes. T. FIJI-based mean pixel intensity quantification of pigmentation phenotype in siNT and siVDAC3 B16 cells in primary human melanocytes (N=4). U. Representative mitochondrial Ca^2+^ imaging traces of CEPIA2mt in siNT control and siVDAC3 B16 cells stimulated with 50µM histamine. V. Quantitation of resting mitochondrial Ca^2+^ with CEPIA2mt in siNT control and siVDAC3 B16 cells where “N” denotes the number of ROIs. W. Quantitation of mitochondrial Ca^2+^ uptake by calculating increase in CEPIA2mt signal (ΔCEPIA2mt) in siNT control and siVDAC3 B16 cells stimulated with 50µM histamine where “N” denotes the number of ROIs. Data presented are mean ± SEM. For statistical analysis, one sample t and Wilcoxon test was performed for panels A, B, C, G, H, I and R, ratio paired t-test was performed for panel Q and T and unpaired Student’s t-test was performed for panel V and W, using GraphPad Prism software. Here, * p<0.05; ** p<0.01; *** p<0.001; and **** p<0.0001. The data underlying panels J, K, L, M, N, O and P are presented in Fig. S3 Data.

VDACs represent the most ubiquitous proteins of the outer mitochondrial membrane, displaying distinct tissue-specific expression patterns [9]. Recent studies report non-redundant and unique functional relevance of each VDAC isoform in cancer cell metabolism, apoptosis, redox metabolism, mitochondrial health and cellular energetics [9,11–13]. Despite the rapid growth of research on VDAC proteins, the VDAC3 isoform remains comparatively under studied and poorly characterized [35]. Therefore, we focused our investigation on VDAC3 to elucidate its functional role and underlying mechanisms in pigmentation biology. We silenced VDAC3 in B16 HD-MSH pigmentation system and observed a significant reduction in VDAC3 protein expression in siVDAC3 transfected cells compared with siNT control (**Fig. S3G and S3H**). As expected, in αMSH-stimulated HD-B16 cells, VDAC3 silencing resulted in hyperpigmentation (**Fig. 3P, 3Q**). Next, we characterized the human VDAC3 siRNA and observed a significant reduction in VDAC3 mRNA levels, while VDAC1 and VDAC2 mRNA levels remained unaffected, in siVDAC3 transfected primary human melanocytes as compared to the siNT control (**Fig. 3R**). We reproducibly recapitulated the hyperpigmentary response upon VDAC3 knockdown in primary human melanocytes as well (**Fig. 3S, 3T**). Collectively, our results clearly demonstrate that VDAC3 negatively modulate pigmentation in mouse and human cellular models. Next, we investigated the role of VDACs in regulating the mitochondrial Ca^2+^ homeostasis of the cells. As expected, we observed that VDAC3 knockdown results in reduced basal mitochondrial Ca^2+^ levels and histamine-stimulated mitochondrial Ca^2+^ uptake (**Fig. 3U-3W**). Similarly, we observed reduced basal mitochondrial Ca^2+^ levels and histamine-stimulated mitochondrial Ca^2+^ uptake in VDAC1 and VDAC2 silenced B16 cells (**Fig. S3I-S3N**).

Here, it is important to note that we observe reduced basal mitochondrial Ca^2+^ levels and mitochondrial Ca^2+^ uptake upon knockdown of Rhot1, VDAC1, VDAC2, and VDAC3, and a concomitant increase in the melanogenic response in all, indicating an inverse relationship between the two. In our previous study, we observed a direct relation between MCU-mediated mitochondrial Ca^2+^ levels and the pigmentary response which was mediated by NFAT2 transcription factor [16]. These observations point towards a dichotomous role of mitochondrial Ca^2+^ dynamics in the regulation of the same process i.e. melanogenesis via unique downstream mechanisms.

### Rhot1 and VDAC3 negatively modulate pigmentation *in vivo*

We next investigated role of Rhot1 and VDAC3 in pigmentation *in vivo* using *Danio rerio* (zebrafish) model system. Zebrafish is a highly relevant model system in pigmentation biology due to the conservation of key physiological pigmentation processes and molecular players across vertebrates [36,37]. Further, transparent embryos and larvae facilitate real-time phenotypic assessment, with melanophores recapitulating key aspects of melanocyte function [23,37]. Thus, zebrafish serves as an excellent *in vivo* model system to assess the role of pigmentation regulators [21,27,32].

To silence the gene of interest, we used a morpholino (MO) based knockdown strategy. MOs are synthetic nucleic acid analogs containing a six-membered morpholine ring instead of canonical deoxyribose/ribose ring, possessing high affinity for RNA and blocking gene expression [38]. The experimental strategy involved incubating zebrafish larvae 24 hours post fertilization (hpf) in phenylthiourea (PTU) containing embryo water for 24h to abrogate pigmentation induction followed by MO injections in the non-pigmented larvae (**Fig. 4A**). The injected embryos were then incubated in normal embryo water for 24h and phenotype was recorded 24 hours post injection (hpi) (**Fig. 4A**).

**Figure 4.**
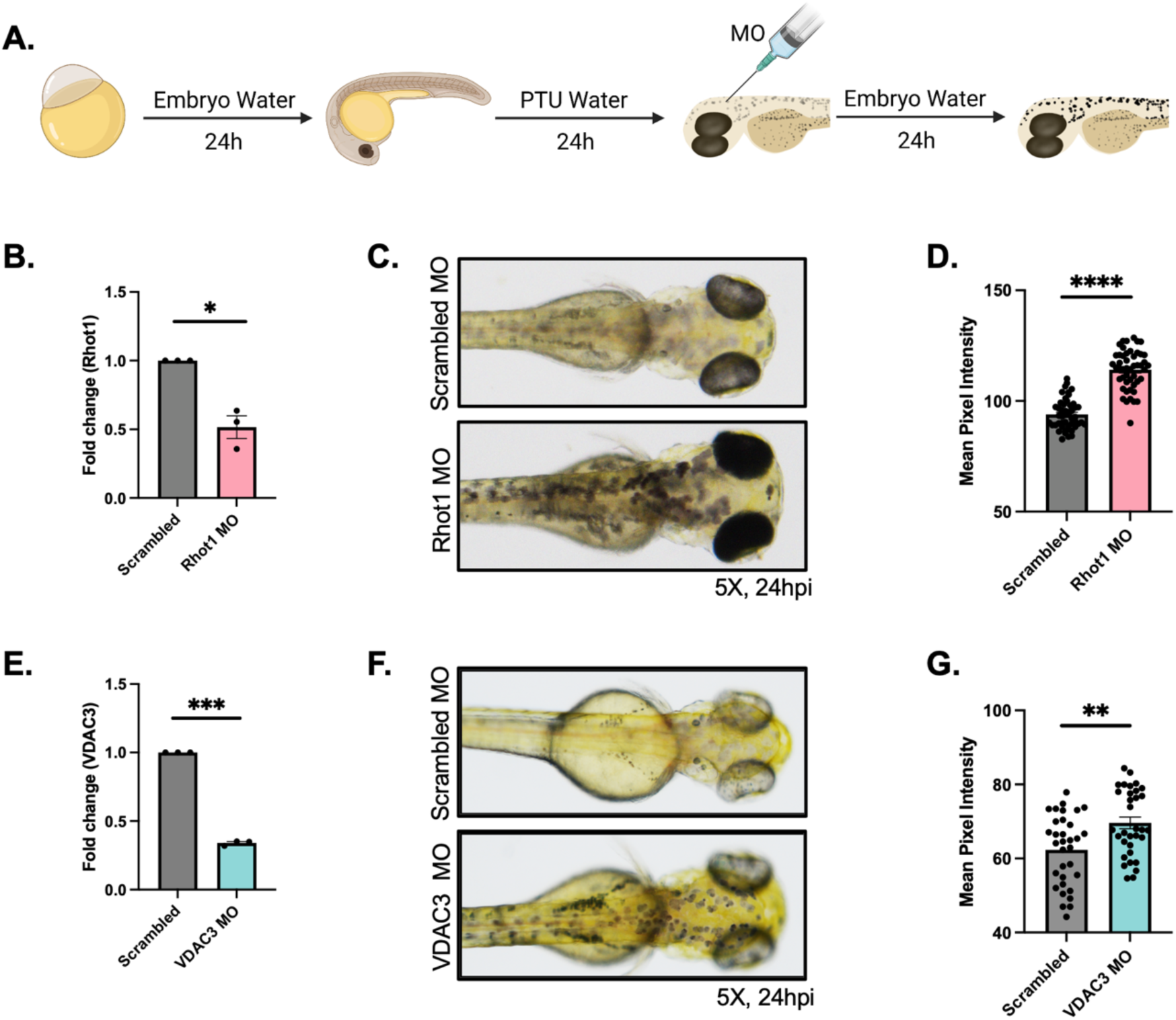
Rhot1 and VDAC3 modulate pigmentation *in vivo*. A. Representative diagram representing the injection strategy for morpholino-based knockdown in zebrafish embryos. B. qRT-PCR analysis confirming morpholino-induced silencing of Rhot1 in zebrafish embryos (N=3). C. Representative bright-field images of wild-type zebrafish larvae injected with either Scrambled control morpholino or morpholino targeting zebrafish Rhot1. D. Mean pixel intensity quantification of pigmentation using FIJI in 72 hpf zebrafish larvae injected with either Scrambled control morpholino or morpholino targeting zebrafish Rhot1 (N=3 independent experiments with 49 embryos/condition, each data point corresponding to one independent larvae/condition). E. qRT-PCR analysis confirming morpholino-induced silencing of VDAC3 in zebrafish embryos (N=3). F. Representative bright-field images of wild-type zebrafish larvae injected with either Scrambled control morpholino or morpholino targeting zebrafish VDAC3. G. Mean pixel intensity quantification of pigmentation using FIJI in 72 hpf zebrafish larvae injected with either Scrambled control morpholino or morpholino targeting zebrafish VDAC3 (N=3 independent experiments with 33 embryos/condition, each data point corresponding to one independent larvae/condition). Data presented are mean ± SEM. For statistical analysis, one sample t and Wilcoxon test was performed for panels B and E, while unpaired Student’s t test was performed for panels D and G, using GraphPad Prism software. Here, * p<0.05; ** p<0.01; *** p<0.001; and **** p<0.0001.

For the first set of experiments, we silenced Rhot1 in zebrafish using Rhot1 MO. We observed around 50% reduction in Rhot1 mRNA expression at 24hpi (**Fig. 4B**). Phenotypically, we observed that Rhot1 knockdown in zebrafish leads to hyperpigmentation as compared to scrambled MO injected larvae, consistent with our *in vitro* experiments (**Fig. 4C**). We quantified the hyperpigmentation phenotype via mean pixel intensity analysis of the embryos on FIJI software (**Fig. 4D**). Next, we used VDAC3 MO to silence VDAC3 in zebrafish. We observed around 70% reduction in VDAC3 mRNA levels at 24hpi (**Fig. 4E**). VDAC3 morphants exhibited a significant increase in black melanophores and were hyperpigmented as compared to scrambled MO injected larvae (**Fig. 4F**). We validated it by carring out mean pixel intensity-based analyses of the larvae to quantify the pigmentary phenotype (**Fig. 4G**). Collectively, our zebrafish data elegantly demonstrates *in vivo* significance of Rhot1 and VDAC3 in pigmentation.

### Unbiased proteomic analysis reveals Kir7.1 functioning downstream of Rhot1 and VDAC3 to modulate pigmentation

To delineate the molecular mechanisms governing pigmentation downstream of Rhot1 and VDAC3, we performed mass spectrometry based global proteomic profiling of B16 cells in the HD-MSH system. αMSH-stimulated siRhot1 and siVDAC3 B16 cells, which clearly exhibited a hyperpigmented phenotype compared to their respective siNT controls, were used (three independent biologicals for each condition) for unbiased proteomics analysis using ZenoTOF 7600 (Sciex) mass spectrometer (**Fig. 5A, 5C**). Sample quality was assessed using standard quality control (QC) metrics and principal component analysis (PCA). QC evaluation indicated high data integrity across samples, while PCA demonstrated tight clustering of biological replicates and clear separation between experimental groups, confirming dataset robustness (**Fig. S4A-S4D**). A 1% false discovery rate cut-off was used at precursor, peptide and protein level for robust identification of protein groups (detailed protocol in Methods section). Post data acquisition, quantification of differential abundance of proteins indicated a robust reduction in Rhot1 and VDAC3 normalized intensity levels in siRhot1 and siVDAC3 conditions, respectively, as compared to their respective siNT controls, thereby validating Rhot1 and VDAC3 silencing in analyzed samples (**Fig. 5B, 5D**). After validating the target protein knockdown in the samples, we analyzed the melanogenic gene expression in test and control conditions. We observed significant increase in GP100 and DCT expression in siRhot1 samples, validating our molecular data (**Fig. 5E, 5F**). Similarly, we also observed increased GP100 and DCT expression upon VDAC3 knockdown (**Fig. 5G, 5H**). Volcano plots of proteomic data demonstrated widespread differential protein expression across conditions. Upon Rhot1 knockdown, 43 proteins appeared to be differentially regulated with a fold change cut-off value ≥1.5 (**Fig. 5I**). Interestingly, in this plot, we observed a significant enrichment (approx. 200% increase) of *Kcnj13* gene which encodes Kir7.1 protein, in Rhot1 silenced condition as compared to siNT control (**Fig. 5I**). Next, volcano plots of siVDAC3 condition vs siNT control exhibited differential regulation of 67 proteins, where we observed some common differentially regulated players as in Rhot1 dataset (**Fig. 5J**). In light of the concordant trends observed across phenotypic, molecular, and ionic flux data, we generated Venn diagram to identify common but differentially regulated candidates across these two independent datasets (**Fig. 5K**). This approach yielded 6 common proteins that were differentially regulated in both datasets (**Fig. 5K**). Excitingly, *Kcnj13* or Kir7.1 emerged as one of the top hits, having extremely high upregulation upon Rhot1 knockdown (∼219%) and VDAC3 knockdown (∼114%) (**Fig. 5L**). It is very interesting because independent proteomic profiling of two distinct mitochondrial Ca^2+^ regulatory proteins resulted in remarkable upregulation of a common ion channel Kir7.1, which is involved in the regulation of cellular K^+^ dynamics. Although we observed five other commonly regulated proteins, we subsequently focused on Kir7.1 as role of K^+^ signaling in skin pigmentation remains unappreciated.

**Figure 5.**
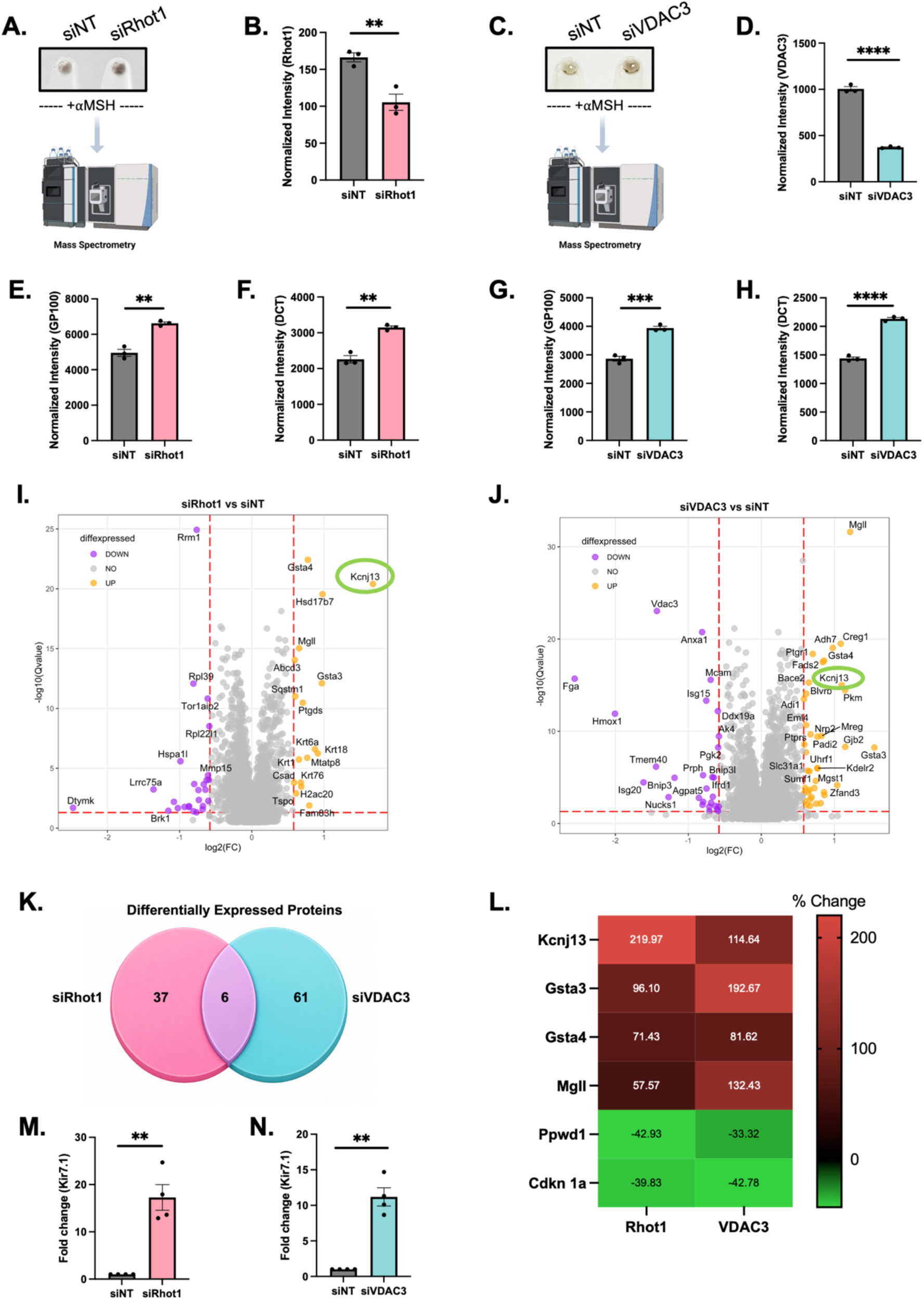
Proteomics analysis identifies Kir7.1 functioning downstream of Rhot1 and VDAC3 to modulate melanogenesis. A. Schematic showing siNT and siRhot1 B16 cells in HD-MSH pigmentation system being subjected to mass spectrometry (MS). B. Validation of Rhot1 knock-down in siNT control vs siRhot1 proteomics samples as indicated by the normalized intensity values obtained from MS analysis (N=3). C. Schematic showing siNT and siVDAC3 B16 cells in HD-MSH pigmentation system being subjected to mass spectrometry. D. Validation of VDAC3 knock-down in siNT control vs siVDAC3 proteomics samples as indicated by the normalized intensity values obtained from MS analysis (N=3). E. Normalized intensity values of GP100 protein in siRhot1 condition compared to siNT control (N=3). F. Normalized intensity values of DCT protein in siRhot1 condition compared to siNT control (N=3). G. Normalized intensity values of GP100 protein in siVDAC3 condition compared to siNT control (N=3). H. Normalized intensity values of DCT protein in siVDAC3 condition compared to siNT control (N=3). I. Volcano plot representing the differentially expressed proteins upon Rhot1 silencing in B16 HD-MSH pigmentation system. J. Volcano plot representing the differentially expressed proteins upon VDAC3 silencing in B16 HD-MSH pigmentation system. K. Venn diagram identifying 6 common differentially expressed proteins in two independent siRhot1 and siVDAC3 proteomic datasets. L. Heatmap representing the expression changes in 6 common differentially expressed proteins in the two proteomic datasets. M. qRT-PCR analysis showing an increase in *Kcnj13* mRNA expression upon Rhot1 silencing in B16 cells (N=4). N. qRT-PCR analysis showing an increase in *Kcnj13* mRNA expression upon VDAC3 silencing in B16 cells (N=4). Data presented are mean ± SEM. For statistical analysis, unpaired Student’s t test was performed for panels B, D, E, F, G and H, while one sample t and Wilcoxon test was performed for panels M and N, using GraphPad Prism software. Here, * p<0.05; ** p<0.01; *** p<0.001; and **** p<0.0001.

Next, we carried out qRT-PCR analysis of Kir7.1 mRNA expression following knockdown of Rhot1 and VDAC3 and observed that silencing Rhot1 or VDAC3 led to significant upregulation of Kir7.1 mRNA expression as well (**Fig. 5M, 5N**). Upon literature survey, we found that Kir7.1 belongs to the family of inward rectifier potassium (Kir) channels that essentially maintain cell membrane potential and potassium (K^+^) fluxes across the cell membrane [39]. Kir7.1 drives a small unitary conductance while establishing the necessary membrane potential in the cells [39,40]. Interestingly, studies in zebrafish have reported Kir7.1 loss-of-function mutations to be responsible for the abnormal movement and distribution of pigment cells in jaguar/obelix mutant fish, which have abnormal stripes patterns [41]. Another study identified evolution of the *Kcnj13* gene (*Kcnj13* gene encodes for Kir7.1 protein) giving rise to divergence in zebrafish pigment pattern formation [42]. Zebrafish *Kcnj13* null mutants display disruption in pigment patterning, implicating a crucial role of *Kcnj13* in color pattern formation in the *Danio* species. *Kcnj13* divergence is responsible for the formation of horizontal and vertical stripe patterns in *Danio rerio* and *Danio aesculapii*, respectively [43]. However, its role in pigmentation biology in other species and the precise molecular mechanism of *Kcnj13* or Kir7.1 mediated zebrafish pigment pattern determination remains elusive. In murine and humans, Kir7.1 has been majorly studied in the retinal pigment epithelium, where it maintains the ionic homeostasis of the subretinal space [44]. Therefore, based on our proteomics data and literature survey, we focused on delineating the role of Kir7.1 in murine and human melanogenesis.

### Kir7.1 modulates melanogenesis positively

Identification of Kir7.1, a K^+^ flux channel as a potential downstream pigment modulator reveals a previously unrecognized link between cellular K^+^ homeostasis and melanogenesis. While the role of cellular Ca^2+^ signaling in melanogenesis is well established, to our knowledge there is no direct evidence that connects cellular K^+^ homeostasis to skin pigmentation. Therefore, we first focused on whether alterations in cellular K^+^ homeostasis can affect melanogenesis. The potassium concentration gradient across the plasma membrane is one of the largest ion gradients in mammalian cells and is essential for maintaining the resting membrane potential, cell volume, and electrical excitability [45,46]. Typically, intracellular K^+^ concentration is in the range of 140-150mM as compared to 3-5mM concentration in the extracellular milieu. We stimulated pigmentation in B16 cells using αMSH and supplemented the cell culture media with increasing concentrations of KCl for 48 hours. KCl treatment disrupts the normal K^+^ gradient across the cell membrane thus affecting the basal K^+^ homeostasis in the cells. We observed increased pigmentation induction with the increasing KCl concentrations (**Fig. 6A, 6B**). For the next set of experiments, we used 2nM valinomycin, a well-known K^+^ ionophore, to extrude cytosolic K^+^ from αMSH -stimulated pigmenting B16 cells. We observed marked visual depigmentation in valinomycin-treated cells compared with DMSO-treated control cells (**Fig. 6C, 6D**). Similarly, valinomycin treatment also induced significant depigmentation in forskolin (Fsk, a chemical inducer of pigmentation by stimulating cAMP generation) stimulated B16 cells as compared to DMSO treated control condition (**Fig. S5A, S5B**). These experiments suggested that K^+^ homeostasis indeed plays a regulatory role in melanogenesis.

**Figure 6.**
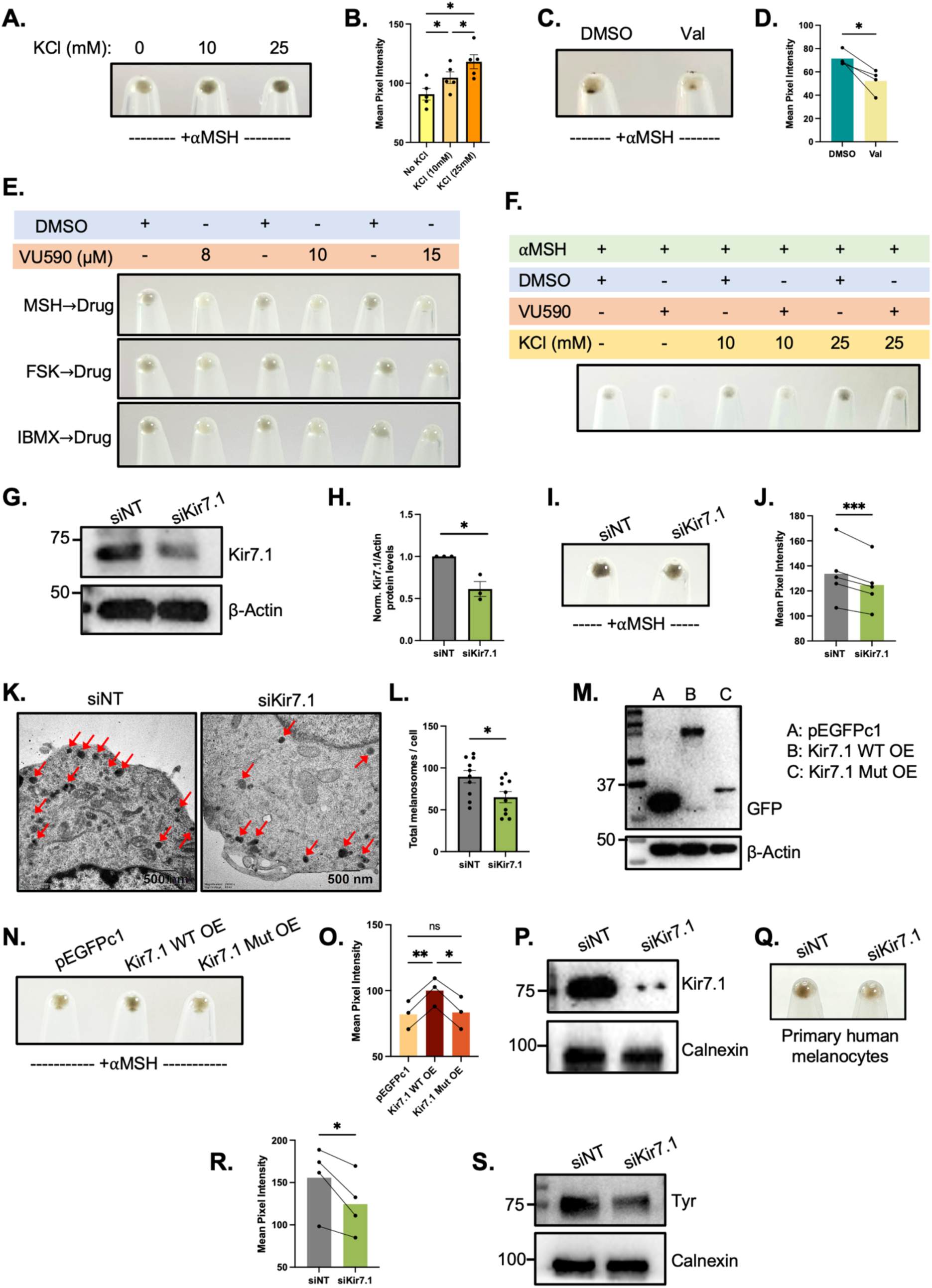
Kir7.1 positively modulates pigmentation. A. Representative pellet picture showing pigmentation phenotype upon increasing concentration of KCl in B16 cell culture media. B. Mean pixel intensity-based pigmentation phenotype quantification upon increasing concentration of KCl in B16 cell culture media (N=5). C. Representative pellet picture showing reduced pigmentation phenotype upon treating B16 cells with Valinomycin as compared to DMSO vehicle control. D. Mean pixel intensity-based pigmentation phenotype quantification upon treating B16 cells with Valinomycin. (N=4). E. Representative pellet picture showing pigmentation phenotype upon treating αMSH, Forskolin or IBMX stimulated B16 cells with increasing concentrations of VU590 (Kir7.1 inhibitor) as compared to DMSO vehicle control. F. Representative pellet picture showing pigmentation phenotype upon combinatorial treatment of increasing KCl concentrations and VU590 on B16 cells stimulated with αMSH. G. Representative western blot validating Kir7.1 knock-down in siKir7.1 condition compared to siNT control in B16 cells (N=3). H. Densitometric quantitation of Kir7.1 protein levels in siKir7.1 condition compared to siNT control (N=3). I. Representative pellet picture comparing pigmentation phenotype of siNT control vs siKir7.1 conditions in HD-MSH pigmentation system. J. Mean pixel intensity-based pigmentation phenotype quantification of siNT and siKir7.1 in HD-MSH pigmentation system (N=5). K. TEM based ultrastructural analysis of siNT control and siKir7.1 B16 cells in HD-MSH pigmentation system (N=10 cells/condition). L. Quantification of total number of melanosomes/cell in siNT control and siKir7.1 B16 cells in HD-MSH pigmentation system (N=10 cells/condition). M. Representative western blot showing GFP-tagged Kir7.1 over-expression upon ectopic expression of WT Kir7.1 and nonsense mutant Kir7.1 in B16 cells. N. Representative pellet pictures comparing pigmentation phenotype of pEGFPc1 control, Kir7.1 WT OE and Kir7.1 Mutant OE in HD-MSH pigmentation system. O. Mean pixel intensity-based pigmentation phenotype quantification of pEGFPc1 control, Kir7.1 WT OE and Kir7.1 Mutant OE in HD-MSH pigmentation system (N=3). P. Representative western blot validating Kir7.1 knock-down in siKir7.1 condition compared to siNT control in primary human melanocytes. Q. Representative pellet pictures comparing pigmentation phenotype of siNT control vs siKir7.1 in primary human melanocytes. R. Mean pixel intensity based pigmentation phenotype quantification of siNT and siKir7.1 conditions in primary human melanocytes (N=4). S. Representative western blot showing tyrosinase levels in siKir7.1 condition compared to siNT control in primary human melanocytes. Data presented are mean ± SEM. For statistical analysis, RM one-way ANOVA was performed for panel B and O, ratio paired t test was performed for panels D, J and R, one sample t and Wilcoxon test was performed for panel H and unpaired Student’s t test was performed for panel L, using GraphPad Prism software. Here, * p<0.05; ** p<0.01; *** p<0.001; and **** p<0.0001. The data underlying panels E, F, P and S are presented in Fig. S5 Data.

Next, we investigated the direct impact of Kir7.1 modulation on pigmentation using its specific pharmacological inhibitor, VU590 [47,48]. We stimulated pigmentation in B16 cells using physiological stimulus αMSH or chemical stimuli Fsk and 3-Isobutyl-1-methylxanthine (IBMX), followed by VU590 treatment at independent doses of 8, 10 and 15µM on the subsequent day. These three stimuli of pigment induction have distinct mechanisms of action. αMSH binds directly to the melanocortin 1 receptor (MC1R), a G-protein coupled receptor (GPCR) present on the melanocyte membrane, triggering cAMP-CREB-MITF cascade leading to pigment synthesis [20]. Forskolin and IBMX bypass the MCIR and lead to pigment induction by increasing intracellular cAMP concentrations by direct adenylyl cyclase (AC) activation and inhibition of phosphodiesterases (PDE), respectively. Upon termination of the experiment i.e. 24 hours post VU590 treatment, we observed significant reduction in pigmentation in the drug-treated cell pellets as compared to the DMSO-treated control pellets (**Fig. 6E**, **S5C**). The observed phenotype was similar for αMSH, Fsk and IBMX-induced pigmentation. Since Fsk and IBMX bypass MC1R, a GPCR, this experiment suggests that the process of pigmentation inhibition by VU590 is a GPCR-independent mechanism. Interestingly, a previous study has also reported G-protein independent function of Kir7.1 in hypothalamic neurons to regulate food intake in rats [47]. Next, we further treated B16 cells with VU590 (8µM) in combination with increasing doses of KCl (0, 10 and 25mM) in presence of αMSH stimulation (**Fig. 6F**).

We observed that specific Kir7.1 inhibition using VU590 almost completely abrogates the KCl induced hyperpigmentation response (**Fig. 6F, S5D**. This suggests that Kir7.1 is playing a crucial role in mediating the KCl induced hyperpigmentation phenotype. Similar observations were made upon treatment of B16 cells with VU590 (8µM) in combination with increasing doses of KCl (0, 10 and 25mM) in presence of Fsk (**Fig. S5E, S5F**).

Further, to functionally validate the potential role of Kir7.1 in melanogenesis, we silenced Kir7.1 in B16 cells and validated the siRNA-induced knockdown via western blotting (**Fig. 6G, 6H**). Kir7.1 silencing in αMSH-stimulated B16 cells clearly resulted in reduced pigmentation phenotype as compared to the siNT control (**Fig. 6I, 6J**). To further visualize the Kir7.1 mediated phenotypic changes, we carried out TEM based ultrastructural analyses where we observed a marked decrease in the total number of melanosomes present per cell in Kir7.1 silenced cells as compared to the control cells (**Fig. 6K, 6L**). We next performed gain of function studies by overexpressing either wild-type (WT) or nonsense mutant isoform of Kir7.1 in B16 cells. Over-expression of human WT Kir7.1 fused with GFP and the nonsense mutant clone with W53X mutation, which has a premature stop codon, was confirmed using western blotting (**Fig. 6M**). We observed that overexpression of WT Kir7.1 resulted in increase in pigmentation, whereas overexpression of the nonsense mutant isoform did not alter the pigmentary response as compared to the control condition (**Fig. 6N, 6O**).

To further substantiate our Kir7.1-mediated phenotypic observations, we extended our findings to another model system, i.e. primary human melanocytes. We silenced Kir7.1 in primary human melanocytes using human Kir7.1 siRNA and observed a significant reduction in Kir7.1 protein expression (**Fig. 6P, S5G**). The knockdown of Kir7.1 in these cells significantly reduced the pigmentation as evident in the pellet pictures and its quantitation (**Fig. 6Q, 6R**).

We also observed a concomitant reduction in the tyrosinase levels upon Kir7.1 inhibition, further strengthening the involvement of Kir7.1 in the melanogenic cascade (**Fig. 6S, S5H**). Collectively, our data reveals that modulation of cellular K^+^ homeostasis plays a crucial role in melanogenesis. Further, we robustly show the critical role of a plasma membrane localized K^+^ flux channel i.e. Kir7.1 in melanogenesis regulation. Our findings from pharmacological inhibition of Kir7.1, loss-and gain-of-function approaches, and validation in primary human melanocytes firmly establish Kir7.1 as a novel positive regulator of melanogenesis.

### Kir7.1 functions downstream of Rhot1 and VDAC3 to modulate melanosome biogenesis

Our data demonstrates that Rhot1 and VDAC3 are negative regulators of pigmentation. Proteomics analyses upon Rhot1 and VDAC3 silencing identified upregulation of Kir7.1 expression in these cells (**Fig. 5**). We show that Kir7.1 is a positive melanogenesis regulator independent of Rhot1 and VDAC3 (**Fig. 6**). We next questioned whether upregulation of Kir7.1 expression downstream of Rhot1 and VDAC3 is essential for mediating the pigmentation response. Therefore, we inhibited the increase in Kir7.1 levels upon Rhot1 knockdown using Kir7.1 siRNA. We estimated the mRNA levels of Rhot1 in the transfected cells and observed a clear reduction in Rhot1 mRNA levels in siRhot1 and siRhot1+siKir7.1 conditions as compared to siNT control (**Fig. S6A**). Next, we quantitated Kir7.1 mRNA levels in transfected cells and observed an increase upon Rhot1 knockdown, consistent with the proteomics data (**Fig. S6B**). In siRhot1+siKir7.1 cells, we observe that siKir7.1 transfection inhibits the increase in Kir7.1 mRNA levels downstream of Rhot1 knockdown, making it similar to that in siNT control cells (**Fig. S6B**). Phenotypically, we observe that while Rhot1 silencing results in hyperpigmentation, inhibition of Kir7.1 along with Rhot1 rescues the pigmentation phenotype (**Fig. 7A-B**). Therefore, Kir7.1 appears to be a necessary mediator of the Rhot1-dependent phenotype.

**Figure 7.**
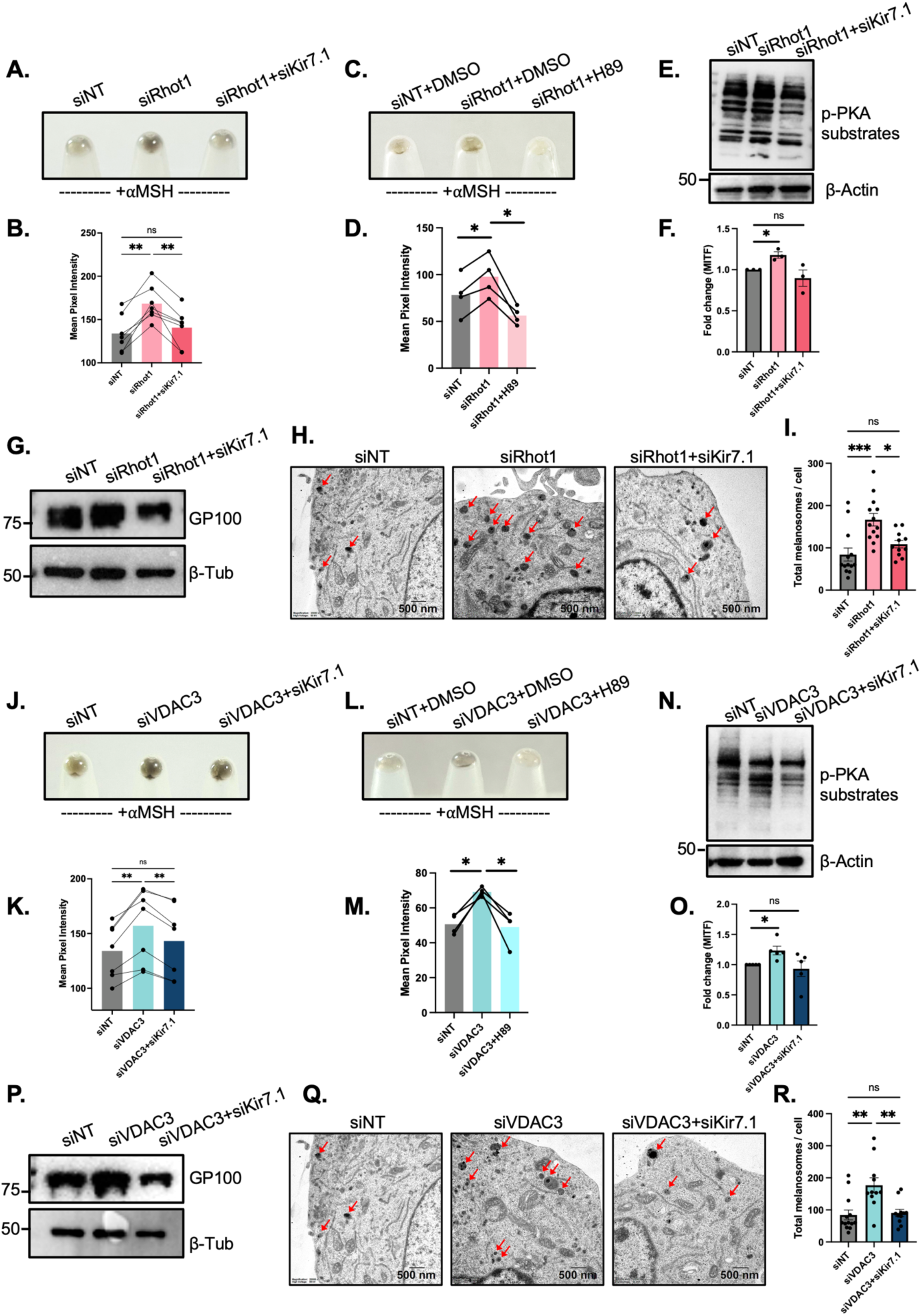
Kir7.1 functions downstream of Rhot1 and VDAC3 to modulate melanosome biogenesis and maturation. A. Representative pellet pictures comparing pigmentation phenotype in siNT control, siRhot1 and siRhot1+siKir7.1 conditions in HD-MSH pigmentation system. B. Mean pixel intensity-based pigmentation phenotype quantification of siNT control, siRhot1 and siRhot1+siKir7.1 conditions in HD-MSH pigmentation system (N=7). C. Representative pellet pictures comparing pigmentation phenotype in siNT control, siRhot1 and siRhot1+H89 conditions in HD-MSH pigmentation system. D. Mean pixel intensity-based pigmentation phenotype quantification in siNT, siRhot1 and H89 treated siRhot1 B16 cells (N=4). E. Representative western blot of cell lysates probed with anti-phospho-PKA substrate (RRXS*/T*) antibody in siNT control, siRhot1 and siRhot1+siKir7.1 conditions in HD-MSH pigmentation system (N=3). F. qRT-PCR analysis of MITF mRNA expression in siNT control, siRhot1 and siRhot1+siKir7.1 conditions in B16 cells (N=3). G. Representative western blot of GP100 levels in siNT control, siRhot1 and siRhot1+siKir7.1 conditions in HD-MSH pigmentation system (N=3). H. TEM based ultrastructural analysis of siNT control, siRhot1 and siRhot1+siKir7.1 conditions in HD-MSH pigmentation system (N=11-13 cells/condition). I. Quantification of total number of melanosomes/cell in siNT control, siRhot1 and siRhot1+siKir7.1 conditions in HD-MSH pigmentation system (N=11-13 cells/condition). J. Representative pellet pictures comparing pigmentation phenotype in siNT control, siVDAC3 and siVDAC3+siKir7.1 conditions in HD-MSH pigmentation system. K. Mean pixel intensity-based pigmentation phenotype quantification of siNT control, siVDAC3 and siVDAC3+siKir7.1 conditions in HD-MSH pigmentation system (N=7). L. Representative pellet pictures comparing pigmentation phenotype in siNT control, siVDAC3 and siVDAC3+H89 conditions in HD-MSH pigmentation system. M. Mean pixel intensity-based pigmentation phenotype quantification in siNT, siVDAC3 and H89 treated siVDAC3 B16 cells (N=4). N. Representative western blot of cell lysates probed with anti-phospho-PKA substrate (RRXS*/T*) antibody in siNT control, siVDAC3 and siVDAC3+siKir7.1 conditions in HD-MSH pigmentation system (N=3). O. qRT-PCR analysis of MITF mRNA expression in siNT control, siVDAC3 and siVDAC3+siKir7.1 conditions in B16 cells (N=5). P. Representative western blot of GP100 levels in siNT control, siVDAC3 and siVDAC3+siKir7.1 conditions in HD-MSH pigmentation system (N=3). Q. TEM based ultrastructural analysis of siNT control, siVDAC3 and siVDAC3+siKir7.1 conditions in HD-MSH pigmentation system (N=11-13 cells/condition). R. Quantification of total number of melanosomes/cell in siNT control, siVDAC3 and siVDAC3+siKir7.1 conditions in the HD-MSH pigmentation system (N=11-13 cells/condition). Data presented are mean ± SEM. For statistical analysis, RM one-way ANOVA was performed for panels B, D, K and M, one sample t and Wilcoxon test was performed for panels F and O, while ordinary one-way ANOVA was performed for panels I and R, using GraphPad Prism software. Here, * p<0.05; ** p<0.01; *** p<0.001; and **** p<0.0001. The data underlying panels C, D, F, K, L and N are presented in Fig. S6 Data.

Literature suggests that αMSH, which induces pigmentation primarily via activating cAMP-Protein Kinase A-MITF axis, also modulates the activity of Kir7.1 [20,23,44,47,48]. So, we questioned if Rhot1 and Kir7.1 mediated melanogenic responses are mediated by this signaling cascade. First, we pharmacologically inhibited Protein Kinase A (PKA), one of the most plausible shared node in the cascade, using H-89. We observed a robust abrogation of pigmentary response in H-89 treated Rhot1 silenced cells as compared to the DMSO treated Rhot1 silenced cells (**Fig. 7C-D**). Further, phosphorylation of PKA substrates was detected using anti-phospho-PKA substrate (RRXS*/T*) antibody, indicating the PKA activity in cells. We observed higher levels of phosphorylated PKA substrates upon Rhot1 knockdown indicating higher PKA activity, which was comparatively reduced in siRhot1+siKir7.1 cells (**Fig. 7E, S6C**). This data is consistent with PKA involvement in mediating the observed pigmentation phenotypes. Further, our qRT-PCR analyses show enhanced MITF mRNA expression upon Rhot1 knockdown, while no changes are observed in MITF expression upon simultaneous knockdown of Rhot1 and Kir7.1 as compared to siNT control condition (**Fig. 7F**) suggesting involvement of MITF in this signaling axis. Next, western blot analysis for MITF target GP100 demonstrated an increase in GP100 levels upon Rhot1 silencing but a reduction upon simultaneous knockdown of Rhot1 and Kir7.1 as compared to siRhot1 condition (**Fig. 7G, S6D**). This data was further substantiated using TEM-based analyses, which clearly show an increase in the total number of melanosomes per cell upon Rhot1 silencing (**Fig. 7H-I**). While upon simultaneous knockdown of Kir7.1 and Rhot1, we observed reduced melanosome biogenesis (**Fig. 7H-I**). Taken together, our data highlights an important role of Kir7.1 in regulating melanogenesis via increased PKA-mediated signaling.

Next, to understand whether upregulation of Kir7.1 expression downstream of VDAC3 is essential for mediating the pigmentation response, we inhibited the increase in Kir7.1 levels upon VDAC3 knockdown using Kir7.1 siRNA. We observed a clear reduction in VDAC3 mRNA levels in siVDAC3 and siVDAC3+siKir7.1 conditions as compared to siNT control (**Fig. S6E**). Next, qRT-PCR analysis revealed an increase in Kir7.1 mRNA levels upon VDAC3 knockdown, consistent with the proteomics data (**Fig. S6F**). In siVDAC3+siKir7.1 cells, we observe that siKir7.1 inhibits the increase in Kir7.1 mRNA levels downstream of VDAC3 knockdown, making it similar to that in siNT control cells (**Fig. S6F**). Phenotypically, we observe that while VDAC3 silencing results in hyperpigmentation while inhibition of Kir7.1 along with VDAC3 rescues the pigmentation phenotype (**Fig. 7J-K**). Next, to understand if VDAC3 mediated melanogenic response is mediated by Protein Kinase A (PKA) signaling, we treated the cells with chemical inhibitor of PKA, H-89. We observed that H-89 treatment in VDAC3 silenced cells inhibits the pigmentation response as compared to the DMSO treated VDAC3 silenced cells (**Fig. 7L-M**). Further, phosphorylation of PKA substrates was detected using western blotting, indicating the PKA activity in cells (**Fig. 7N, S6G**). We observed increased levels of phosphorylated PKA substrates upon VDAC3 knockdown, which were comparatively reduced in siVDAC3+siKir7.1 cells (**Fig. 7N, S6G**). This data suggests PKA mediates the observed pigmentation phenotypes. Consistent with our phenotypic observations, we observed enhanced MITF mRNA expression upon VDAC3 silencing and no changes in MITF expression upon simultaneous knockdown of VDAC3 and Kir7.1 as compared to the siNT control (**Fig. 7O**). Western blot data exhibits that co-silencing of Kir7.1 and VDAC3 inhibits the increase in GP100 expression as observed in siVDAC3 condition (**Fig. 7P, S6H**). Finally, we carried out TEM studies to investigate melanosome biogenesis, which suggest an increase in the total number of melanosomes per cell upon VDAC3 silencing compared to siNT control **(Fig. 7Q-R).** While when Kir7.1 and VDAC3 are silenced simultaneously the number of melanosomes per cell are rescued, which are comparable to the siNT control **(Fig. 7Q-R).** Taken together, our data demonstrates that increase in Kir7.1 expression downstream of Rhot1 and VDAC3 is an essential determinant of the pigmentation phenotype and inhibiting its upregulation downstream of Rhot1 and VDAC3 abrogates the hyperpigmentary phenotype by curtailing the PKA-MITF axis mediated melanogenesis.

### Kir7.1 modulates cellular K^+^ homeostasis downstream of Rhot1 and VDAC3

To have a mechanistic understanding of Kir7.1 channel function during pigmentation, we investigated its role in the regulation of cellular K^+^ levels and the resultant changes in cell membrane potential-both of which are key processes regulated by Kir7.1 channel. To visualize cellular K^+^ in live cells, we used ION Potassium Geen-2 AM dye. We firstly validated its ability to measure changes in cellular K^+^ levels by staining DMSO and Valinomycin treated B16 cells. Valinomycin is a K^+^ ionophore, therefore we observed a stark reduction in the green fluorescence of the K^+^ dye upon acute drug treatment as compared to the DMSO control (**Fig.S7A**). Subsequently, we questioned whether resultant or induced alterations in Kir7.1 expression downstream of Rhot1 and VDAC3 can modify cellular K^+^ levels and membrane polarization. Interestingly, we observed that Rhot1 knockdown, which leads to increased expression of Kir7.1 downstream, results in intracellular K^+^ accumulation, whereas silencing Kir7.1 in combination with Rhot1 silencing rescues the K^+^ accumulation in the cells (**Fig. 8A, 8B**). Further, K^+^ accumulation results in a concomitant depolarization in siRhot1 cells as indicated by increased DiBAC4(3) (Bis-(1,3-Dibutylbarbituric Acid Trimethine Oxonol) dye signal, which is rescued when Kir7.1 is silenced in combination with Rhot1 (**Fig. 8C, 8D**). We used KCl treatment to induce depolarisation in B16 cells as a positive control to establish the dye functioning (**Fig. S7B, S7C**). Similar to our observations with Rhot1 and Kir7.1, VDAC3 silencing in B16 cells, which results in upregulation of Kir7.1 expression, leads to K^+^ accumulation in the cells, whereas Kir7.1+VDAC3 knockdown reduces the K^+^ accumulation in the cells as compared to the VDAC3 silenced cells (**Fig. 8E, 8F**). Concomitantly, we observed depolarization upon VDAC3 knockdown, which was rescued upon combination knockdown of Kir7.1 and VDAC3 (**Fig. 8G, 8H**).

**Figure 8.**
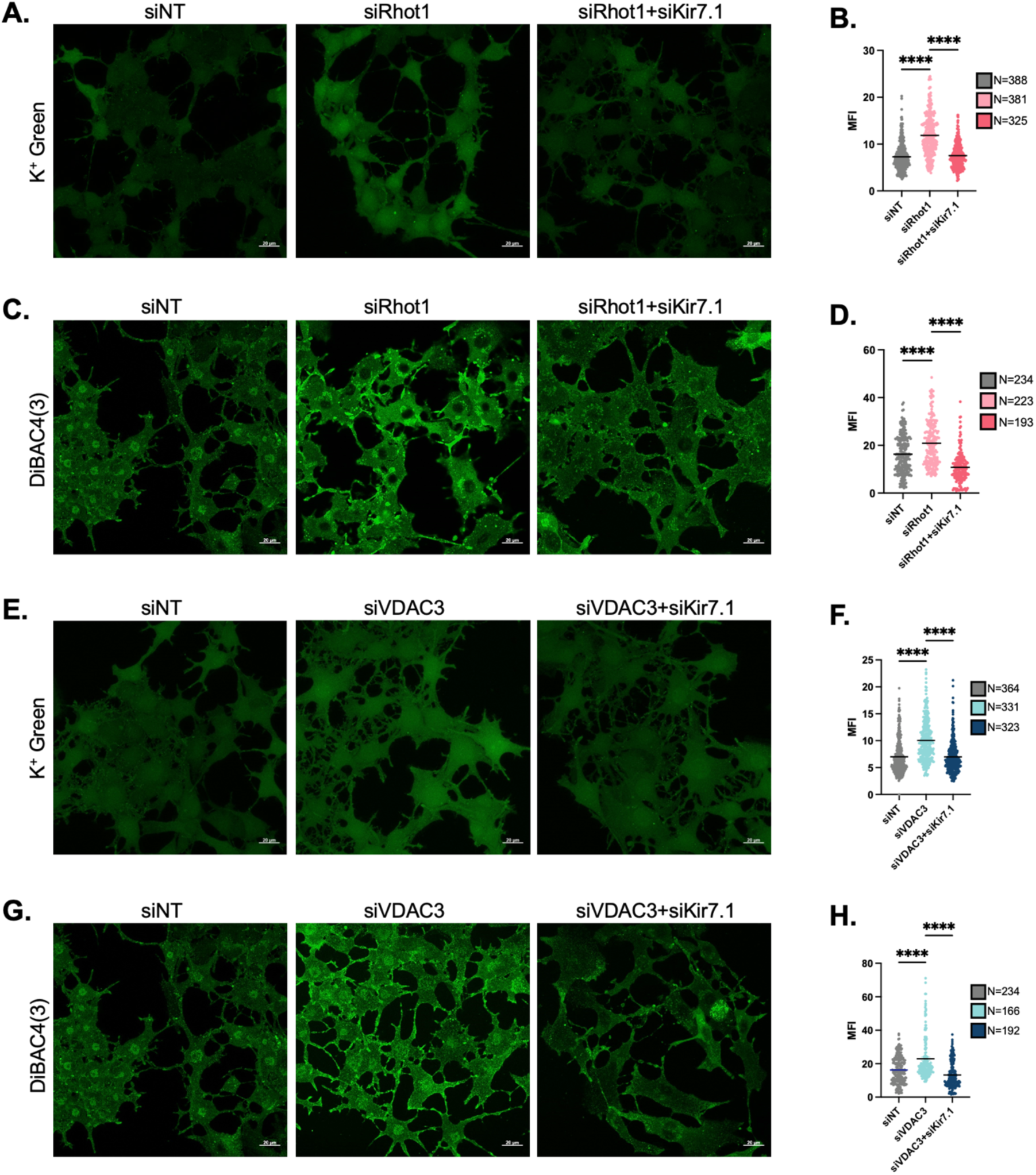
Kir7.1 affects cellular K^+^ homeostasis and membrane potential downstream of Rhot1 and VDAC3. A. Representative images for cellular K^+^ levels in siRhot1 and siRhot1+siKir7.1 B16 cells compared to siNT control (Scale: 20μm). B. Quantification of mean fluorescence intensity indicating cellular K^+^ levels in siRhot1 and siRhot1+siKir7.1 B16 cells compared to siNT control, where ‘N’ denotes the number of cells imaged. C. Representative images for dye-based cell membrane potential confocal imaging in siRhot1 and siRhot1+siKir7.1 B16 cells compared to siNT control (Scale: 20μm). D. Quantification of mean fluorescence intensity indicating cell membrane potential in siRhot1 and siRhot1+siKir7.1 B16 cells compared to siNT control, where ‘N’ denotes the number of cells imaged. E. Representative images for cellular K^+^ levels in siVDAC3 and siVDAC3+siKir7.1 B16 cells compared to siNT control (Scale: 20μm). F. Quantification of mean fluorescence intensity indicating cellular K^+^ levels in siVDAC3 and siVDAC3+siKir7.1 B16 cells compared to siNT control, where ‘N’ denotes the number of cells imaged. G. Representative images for dye-based cell membrane potential confocal imaging in siRhot1 and siVDAC3+siKir7.1 B16 cells compared to siNT control (Scale: 20μm). H. Quantification of mean fluorescence intensity indicating cell membrane potential in siRhot1 and siVDAC3+siKir7.1 B16 cells compared to siNT control, where ‘N’ denotes the number of cells imaged. Data presented are mean ± SEM. For statistical analysis, ordinary one-way ANOVA was performed for panels B, D, F and H, using GraphPad Prism software. Here, * p<0.05; ** p<0.01; *** p<0.001; and **** p<0.0001.

Interestingly, we observed that pharmacological blocking of Kir7.1 with VU590 increased K^+^ accumulation in the cell (**Fig. S7D, S7E**). As a result of increased intracellular K^+^, the cells treated with VU590 exhibit a significant depolarization (**Fig. S7F, S7G**). Our pharmacological data is well supported by siRNA-mediated genetic knockdown data, where we observed a similar intracellular K^+^ accumulation in Kir7.1 silenced cells as compared to the siNT control (**Fig. S7H, S7I**). Consequently, Kir7.1 silencing also increased the DiBAC4(3) fluorescence signal indicating cellular depolarization (**Fig. S7J, S7K**). Based on our data, since Kir7.1 expression increases upon Rhot1 as well as VDAC3 silencing, the channel K^+^ efflux function could be blocked as a feedback mechanism, resulting in the observed K^+^ accumulation in the cells and depolarization.

Overall, this data hints towards a dichotomous regulation of pigmentation and cellular K^+^ homeostasis by Kir7.1 modulation-where increased Kir7.1 expression downstream of Rhot1 and VDAC3 is essential for hyperpigmentation phenotype, K^+^ accumulation in the cell and cellular depolarization, all of which are rescued when we inhibit the increase of Kir7.1 levels downstream Rhot1 and VDAC3 knockdown. But direct inhibition or silencing of the Kir7.1 results in reduced pigmentation, K^+^ accumulation and cellular depolarization. This observation emphasizes the importance of the initiation hub of the signal-indirect Kir7.1 modulation downstream of Rhot1 or VDAC3 knockdown, and in contrast, direct modulation of Kir7.1 using inhibitor or siRNA-both of which are depolarizing stimuli leading to cellular K^+^ accumulation, but induce different pigmentary responses.

### MITF regulates Kir7.1 expression in the pigmentation cascade

Since our data clearly show that Kir7.1 mRNA and protein expression are enhanced upon silencing Rhot1 or VDAC3, we began investigating the molecular details underlying thisregulation. To delineate the transcriptional regulation of Kir7.1 in melanogenesis, we conducted extensive bioinformatics analysis of the mouse Kir7.1 promoter using three independent tools: Eukaryotic Promoter Database (EPD), ContraV3 and PSCAN [49–52]. The computational analyses revealed potential MITF binding sites on the Kir7.1 promoter (**Fig. 9A**). PSCAN was used to identify putative transcription factor-binding sites within the Kir7.1 promoter using JASPAR Core 2020 nonredundant transcription factor position weight matrices (**Fig. 9B**). We also analysed the Kir7.1 promoter for MITF binding sites using the EPD-Search Motif Tool. Mouse Kir7.1 promoter sequence was retrieved and analysed for MITF binding sites, applying a stringent *p*-value cut-off of *p* = 0.01 (**Fig. 9C**). Next, we performed cross-species alignment of the conserved MITF binding sites on Kir7.1 promoter using the ContraV3 transcription factor binding analysis tool, applying a core similarity threshold of 0.95 and a matrix similarity threshold of 0.85 (**Fig. 9D**). The predicted TF-binding sites are evolutionarily conserved across diverse vertebrate taxa, including reptiles, birds, and multiple mammalian species. Importantly, all three tools, using independent algorithms, consistently identified several MITF binding sites on the Kir7.1 promoter. Thus, our stringent computational analyses suggest that MITF transcription factor may transcriptionally regulate Kir7.1 to modulate melanogenesis.

**Figure 9.**
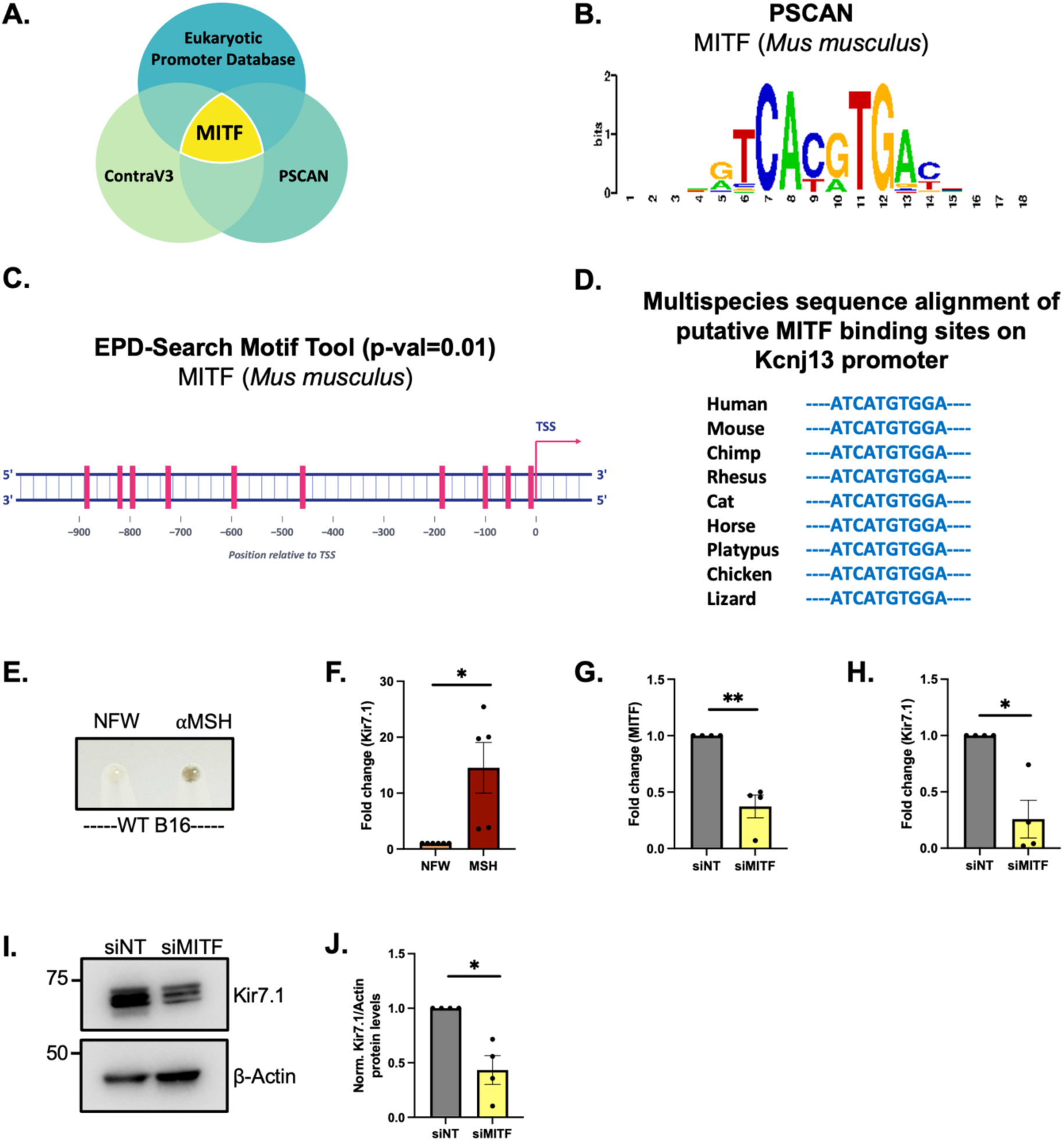
MITF regulates Kir7.1 expression during melanogenesis. A. In-silico analysis via Eukaryotic Promoter Database, ContraV3 and P-SCAN identifies MITF as a potential transcriptional regulator of Kir7.1. B. Identification of putative MITF binding sites on the Mouse *Kcnj13* promoter using the EPD-Search Motif Tool at P-value cut-off of 0.01 C. Bioinformatic characterization of conserved MITF binding sites on the cross-species alignment of the *Kcnj13* promoter using the ContraV3 transcription factor binding analysis tool. D. Position weight matrix of mouse MITF consensus binding sequence obtained via PSCAN. E. Representative pellet picture comparing pigmentation phenotype in WT B16 cells upon treatment with NFW (control) and αMSH. F. qRT-PCR based Kir7.1 mRNA levels quantitation in NFW treated non-pigmented and αMSH treated pigmented B16 cells (N=5). G. qRT-PCR analysis confirming MITF knockdown in siMITF B16 cells as compared to siNT control (N=4). H. qRT-PCR based Kir7.1 mRNA levels quantitation in siMITF cells compared to siNT control (N=4). I. Representative western blot of Kir7.1 levels in siMITF B16 cells as compared to siNT control (N=4). J. Densitometric quantitation showing Kir7.1 levels in siMITF B16 cells compared to siNT control (N=4). Data presented are mean ± SEM. For statistical analysis, one sample t and Wilcoxon test was performed for panels F, G, H and J, using GraphPad Prism software. Here, * p<0.05; ** p<0.01; *** p<0.001; and **** p<0.0001.

Initially, to understand the transcriptional regulation of Kir7.1 in melanogenesis downstream of MITF, we treated the B16 cells with NFW and αMSH as αMSH activates MITF transcription program and thereby induces pigmentation (**Fig. 9E**). αMSH is a well-known inducer of the MITF-mediated melanogenic cascade. We next compared the expression of Kir7.1 in these cells and observed higher expression of Kir7.1 in αMSH-treated pigmented cells as compared to the NFW-treated non-pigmented control (**Fig. 9F**). Further, we silenced MITF in B16 cells and validated the MITF knockdown via qRT-PCR analysis (**Fig. 9G**). We observed that upon MITF silencing, Kir7.1 mRNA expression is significantly reduced (**Fig. 9H**). Further, we observed a similar reduction in Kir7.1 protein expression as well (**Fig 9I, 9J**). These data show a critical role of MITF in regulating Kir7.1 expression during melanogenesis.

### Kir7.1 is differentially expressed in melanogenic disorders

Our study extensively demonstrates that Kir7.1 is a critical component regulating melanogenesis-both independently and downstream of other melanogenic effectors such as Rhot1 and VDAC3. This establishes Kir7.1 as a novel critical regulatory protein that is expressed downstream of MITF, functions to alter melanocyte K^+^ homeostasis and melanogenesis. We next investigated its involvement in hypopigmentary pathologies such as vitiligo and hyperpigmentary disorders such as melasma. Using the publicly accessible microarray datasets, we analysed the skin gene expression levels of *Kcnj13* in control and test groups. We observed that in vitiligo, where the lesional skin exhibits marked depigmentation and perilesional skin exhibits hypopigmentation, *Kcnj13* expression is significantly reduced in the lesional depigmented skin as compared to the peri-lesional and non-lesional skin, which exhibit higher pigmentation (**Fig. 10A**, Database ID: GSE65127). In an independent dataset, we similarly observed a significant reduction in *Kcnj13* expression in the vitiligo skin as compared to the control skin samples (**Fig. 10B**, Database ID: GSE53146). Interestingly, we observed an opposite trend in melasma-affected skin samples, where *Kcnj13* expression was markedly increased in hyperpigmented melasma lesions as compared to the non-lesional skin (**Fig. 10C**, Database ID: GSE72140).

**Figure 10.**
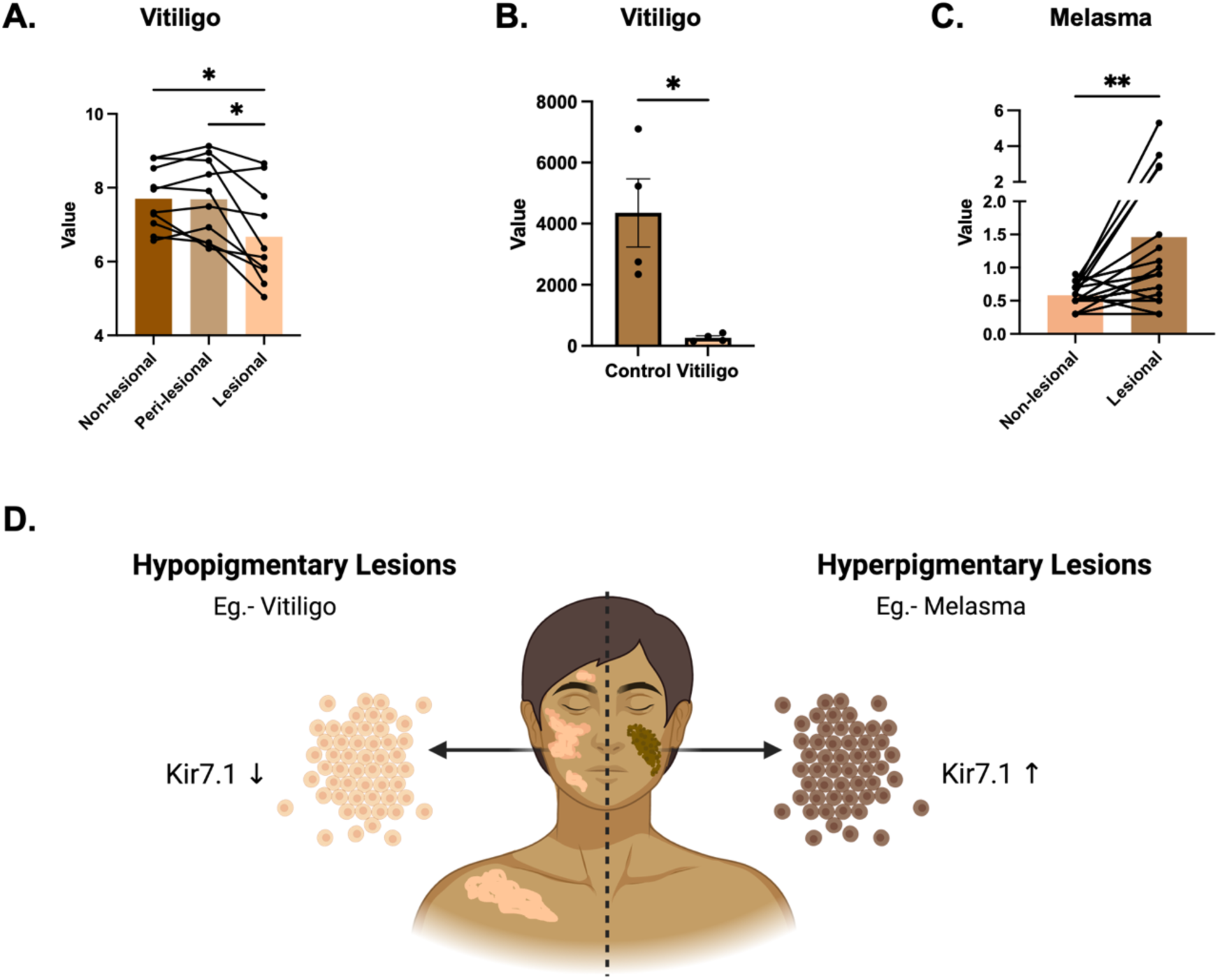
Differential expression of Kir7.1 in melanogenic disorders. A. Differential Kir7.1 expression in non-lesional, peri-lesional and lesional vitiligo skin from publically available database (ID: GSE65127; N=10) B. Kir7.1 expression in control vs vitiligo skin from publically available database (ID: GSE53146; N=4) C. Kir7.1 expression in non-lesional vs lesional skin in melasma paitents from publically available database (ID: GSE72140; N=17) D. Schematic showing differential expression of Kir7.1 in hypopigmentary vs hyperpigmentary lesions of melanogenic disorders. Data presented are mean ± SEM. For statistical analysis, RM one-way ANOVA was performed for panel A, Student’s t test was performed for panels B and ratio paired t test was performed for panel C, using GraphPad Prism software. Here, * p<0.05; ** p<0.01; *** p<0.001; and **** p<0.0001.

Overall, analyses from three independent microarray datasets of hypo-and hyper-pigmentary pathologies vitiligo and melasma, respectively, indicate a positive correlation of *Kcnj13* gene expression with the pigmentation of the affected lesional skin (**Fig. 10D**). This data, along with other data presented in this study, suggests a possible role for Kir7.1 in pigmentary disease pathogenesis, as well as its potential as a biomarker and a candidate therapeutic target for pigmentary disease management.

## Discussion

Cellular ionic signaling plays a key role in regulating melanogenesis and skin pigmentation. Complex ion transport mechanisms across the membranes of various cellular organelles, such as mitochondria, ER, melanosomes as well as plasma membrane, critically modulate melanin synthesis, pigment storage and melanosome transfer [53]. Alterations in ion homeostasis exert their downstream effects by modulating enzyme activity, activating transcription factors, regulating organelle pH, and affecting other processes. The role of Ca^2+^ in pigmentation has been established by several studies [16,21,31,32,54,55]. Our group has highlighted the significance of STIM1, an ER Ca^2+^ sensor, and its interaction with the plasma membrane resident adenylyl cyclase 6 (ADCY6) in governing melanogenesis upon αMSH-mediated ER Ca^2+^ depletion [23]. This leads to cAMP generation and forms a positive feedback loop that sustains cAMP levels essential for induction of pigmentation genes. It is intriguing that while mitochondrial Ca²⁺ dynamics are well known to regulate diverse physiological phenomena, their role in melanosome biology and melanogenesis has remained largely unexplored. We recently revealed a critical role for IMM-localised MCU-mediated mitochondrial Ca^2+^ uptake in positively modulating melanogenesis using cellular, zebrafish, and transgenic mouse model systems [16]. Based on our observations that mitochondrial Ca^2+^ dynamics show distinct trends with pigmentation induction and modulation, we robustly screened the key mitochondrial Ca^2+^ handling machinery proteins to identify novel modulators of pigmentation and dissect the underlying molecular choreography that contributes towards the pigmentary phenotype.

In this study, using unbiased siRNA-based screening of the mitochondrial Ca^2+^ handling proteins, we identified six novel mitochondria-localized melanogenesis modulators namely, Rhot1, VDAC1, VDAC2, VDAC3, SLC25a23 and SLC25a24 **(Fig. 1)**. Here, we focused on Rhot1 and VDACs (**Fig. 2 and 3**). Using four independent models of pigmentation-B16 LD pigmentation model, αMSH-stimulated pigmentation in B16 cells and primary human melanocytes and *in vivo* zebrafish pigmentation model, we establish a critical role of Rhot1 and VDACs (VDAC1/2/3) as negative modulators of pigmentation (**Fig. 2 and 3**). These data are well supported by increased expression of the key melanogenic proteins such as DCT, tyrosinase, GP100 or increased tyrosinase activity upon the target protein silencing (**Fig. 2 and 3**). Two recent reports identified VDAC1 as a negative regulator of pigmentation [33,34] . They show that VDAC1 depletion in MNT-1 and Melan-a cells elevates cytosolic Ca²⁺, activating calcineurin-dependent CRTC1 dephosphorylation and nuclear translocation, which drives MITF transcription and melanogenic gene expression [33,34]. However, in literature, there is no report of VDAC2 and VDAC3 in the modulation of melanogenesis. VDAC3, albeit sharing 60-70% sequence identity with its paralogs, has several distinct features contributing to its unique functions, yet it remains the least understood isoform [9,12,35]. Although the VDAC isoforms are ubiquitously expressed, VDAC3 is especially abundantly expressed in cells and tissues of the germinal lineage. VDAC3 has been implicated in various pathologies like cancer, viral infections, cerebral malaria and chronic unpredictable stress (CUS) [35]. Our study now highlights the critical role of the least studied VDAC isoform-VDAC3, in orchestrating the melanogenic response (**Fig. 3**). We further extended our phenotypic findings in independent cellular models to an *in vivo* model, i.e., zebrafish, where we observed a compelling hyperpigmentation response in Rhot1 and VDAC3 morphants, highlighting the physiological relevance of these proteins in vertebrate pigmentation (**Fig. 4**).

Intriguingly, we observed that despite the knockdown of two different mitochondrial OMM proteins, *Kcnj13,* which encodes Kir7.1, a plasma membrane localized K^+^ flux channel, emerged as one of the most prominently upregulated proteins in the two independent datasets (**Fig 5**). Interestingly, Kir7.1 determines pigment pattern formation in zebrafish, but mechanistic details remain unknown [41–43]. Genetic channelopathies associated with Kir7.1 loss-of-function include Snowflake Vitreoretinal Degeneration (SVD) and Leber Congenital Amaurosis (LCA), both of which affect retinal function and vision [42]. More recently, it has been shown to play an important role in the regulation of food intake in mice, uteroplacental function, smooth muscle cytoskeletal organization for mouse trachea formation and determining oligodendrocyte integrity in the mouse optic nerve [43–48]. However, its role in mammalian pigmentation and melanogenesis remains unexplored. Upon discovering the involvement of a K^+^ handling channel in melanogenesis, we first established that cellular K^+^ homeostasis is crucial for modulating the melanogenic outcome in mammalian models (**Fig. 6**). Thereafter, we demonstrated that Kir7.1, independently, is a positive modulator of pigmentation in B16 and human cellular models using pharmacological inhibition of the channel, siRNA-mediated gene silencing and overexpression studies (**Fig. 6**). Ghamari- Langroudi *et al*. have previously reported that Kir7.1 couples to MC4R in G-protein independent manner in hypothalamic neurons, where they also observed a similar coupling of MC1R with Kir7.1 [47]. Interestingly, both neural cell types and melanocytes are derived from the neural crest cells originated in the ectoderm but MC1R, a paralog of MC4R, is the major melanocortin receptor expressed in melanocytes [56]. In parallel to this study, we observed G- protein independent function of Kir7.1 in melanocytes **(Fig. 6)**. Pharmacological inhibition of Kir7.1 using VU590 abrogated the αMSH-induced pigmentation. It also decreased the pigmentation induced by Forskolin and IBMX that bypass MC1R binding and directly elevate cAMP levels. To the best of our knowledge, this is the first report on cellular K^+^ dynamics and K^+^ handling protein, Kir7.1 regulating melanogenesis.

Further, combined knockdown of Rhot1 or VDAC3 with Kir7.1 prevented upregulation of Kir7.1 and also abolished the hyperpigmentary response. This highlights the significance of Kir7.1 upregulation downstream of Rhot1 and VDAC3 silencing in mediating the hyperpigmentary response (**Fig. 7**). Concomitant changes were observed in melanosome biogenesis and maturation as well. We also show that Kir7.1 modulation during melanogenesis significantly alters the cellular K^+^ accumulation and cell membrane potential (**Fig. 8**). Recently it has been shown that Opsin3, previously described to negatively regulate pigmentation in melanocytes, also regulates MC4R–Kir7.1 signaling in hypothalamic neurons to promote food intake in mice [48,57]. The authors observed Opsin3 silencing enhances Kir7.1 expression, but reduces the K^+^ flux through the channels [48]. Notably, our results closely parallel with this study as Rhot1 and VDAC3 negatively modulate pigmentation by upregulating Kir7.1 expression. Despite enhanced Kir7.1 expression upon Rhot1 and VDAC3 silencing, cytosolic K^+^ levels remain high, indicating diminished K^+^ flux through Kir7.1. It is therefore noteworthy that some of the Kir7.1 channel signaling signatures overlap in neuronal and melanocyte homeostasis and point towards conserved regulatory mechanisms.

We next delved into understanding the molecular basis of high Kir7.1 upregulation during Rhot1/VDAC3-mediated melanogenesis **(Fig. 9)**. Our data suggests shows that Kir7.1 expression is increased upon physiological melanogenic stimulation with αMSH which activates melanogenesis by triggering the PKA-CREB-MITF signaling axis (**Fig. 9**). We further observed that Kir7.1 mRNA and protein expression are diminished upon MITF silencing in B16 cells. This reveals that melanocytes have adopted a signaling cascade, involving MITF transcription factor, to augment Kir7.1 expression during αMSH-stimulated melanogenesis. Earlier literature also links cellular K^+^ handling machinery and its regulation by MITF transcription factor. One study highlights the role of MITF in maintaining the neuroplasticity of olfactory bulb (OB) projection neurons by regulation the *Kcnd3* genes, which encode the voltage-gated K^+^ channel Kv4.3 [58]. Transcriptome analysis of cochlear stria vascularis in *Mitf-m* knockout mice having severe hearing disability indicated downregulation of genes of the melanogenic cascade and inward rectifier K^+^ channels *Kcnj10* and *Kcnj13* [59]. Bulk RNA-seq analysis of *Mitf* knockout *Xenopus tropicalis* dorsal and ventral skin samples also showed reduced *Kcnj13* gene expression [60]. However, the understanding of the molecular mechanism connecting the *Kcnj13* gene expression and MITF still remains elusive. In this study we show the regulation of *Kcnj13* gene expression downstream of MITF through extensive bioinformatics analyses coupled with molecular assays (**Fig. 9**).

Another intriguing observation of this study is a tight association between Kir7.1 expression and the pigmentary disorders (**Fig. 10**). Analysis of Kir7.1 expression in three independent publicly available skin transcriptomics datasets from patients with vitiligo and melasma revealed that Kir7.1 expression is directly proportional to pigmentation in diseased lesions. Interestingly, an earlier study reported markedly distinct kinetics of MC4R-mediated cAMP signaling and Kir7.1 regulation [47]. As the cAMP response diminished rapidly after signal induction, Kir7.1 regulation progressively increased and persisted for longer durations following α-MSH stimulation [47] .These findings highlight the ability of GPCRs to engage parallel signaling mechanisms to regulate ion channel activity, independently of their conventional G-protein-mediated pathways, and underscore the importance of receptor–ion channel interactions in shaping cellular responses. Similarly, in melanocytes, robust upregulation of Kir7.1 during pigmentation could be essential to maintain a feed-forward loop that ensures sustained melanogenesis after transient pigment stimulus such as cAMP. This represents a potential confounding factor that maintains increased expression of Kir7.1 and thereby high melanin synthesis in hyper-pigmentary disorders like melasma. These observations suggest that Kir7.1 could be an important mediator of pigmentary disease outcomes and could be targeted to manage melanogenic anomalies in the skin. Going forward, it would be worth investigating its therapeutic or biomarker potential for the management of pigmentary disorders. To date, there are no FDA-approved drugs targeting this channel. Future studies aimed at investigating the draggability of Kir7.1 to alleviate pigmentary disorders could hold considerable promise for transforming current therapeutic strategies and advancing the treatment landscape.

Taken together, this study identifies novel modulators of pigmentation, namely-Rhot1, VDAC1, VDAC2, VDAC3 and Kir7.1. Further, we reveal the importance of K^+^ ion signaling in pigmentation and report a novel Rhot1/VDAC3-PKA-MITF-Kir7.1 axis contributing to pigmentation (**Fig. 11**). The classical αMSH-mediated pigmentation pathway requires MITF-mediated transcriptional upregulation of Kir7.1 to elicit the pigmentation responses. Collectively, our findings position K^+^ ion signaling as a previously unrecognised regulator of melanogenesis, and our work opens a new frontier for research with potential implications for both fundamental pigment cell biology and the development of therapeutic strategies for pigmentary disorders. The implications of such signaling modules may extend well beyond melanogenesis, such as in neuronal disorders, in other physiological and pathological contexts. Future studies exploring this axis in other biological contexts represents an exciting avenue for future research and may provide novel insights into the regulation of cellular function and disease.

**Figure 11.**
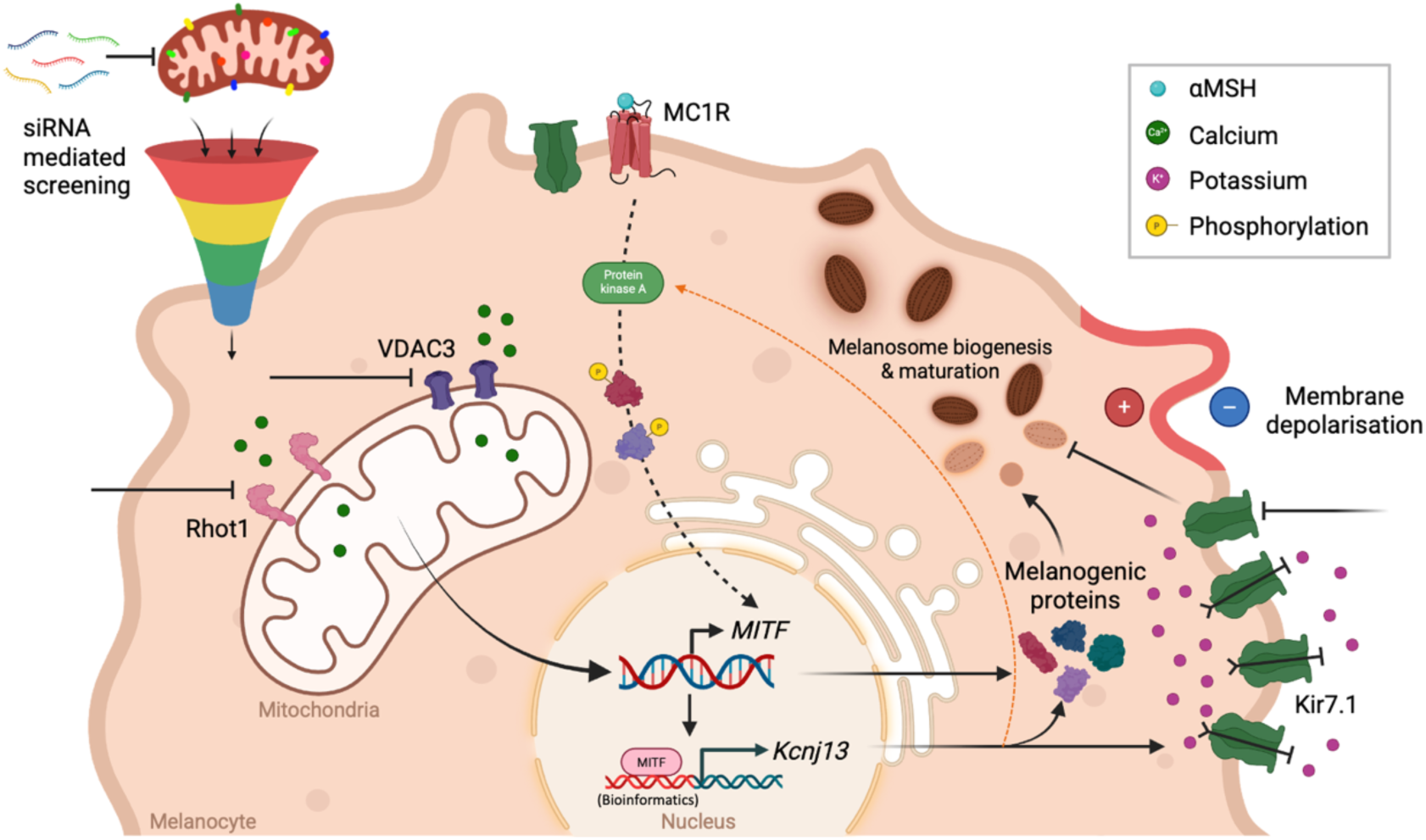
Rhot1 and VDAC3 induced MITF-mediated transcriptional regulation of Kir7.1 orchestrates pigmentation. Graphical summary of the study illustrating the unbiased siRNA-based screening mediated identification of novel mitochondria-localized modulators of pigmentation-Rhot1 and VDAC3. Rhot1 and VDAC3 silencing induce PKA activation and MITF expression, which results in *Kcnj13* upregulation. *Kcnj13* encoded Kir7.1 protein alters cellular K^+^ homeostasis, cell membrane potential and augments melanosome biogenesis and maturation, thus, contributing to the hyperpigmentary response. (Created using Biorender.com)

## Acknowledgements

This work was supported by the DBT/Wellcome Trust India Alliance Fellowship (IA/I/19/2/504651) to RKM. RKM also acknowledges funding support from RCB Institutional Core funding (R25226). The authors thank the members of the Motiani laboratory for the critical discussions. KA acknowledges her Junior and Senior Research Fellowship from DBT, India. IP and AJ acknowledge their Junior Research Fellowship from CSIR, India. AT acknowledges his Junior Research Fellowship from DBT, India. AB acknowledges his Junior and Senior Research Fellowship from DBT, India.

## Author contributions

Kriti Ahuja: Formal analysis; Investigation; Visualization; Methodology; Writing-original draft. Ishita Panthri: Formal analysis; Investigation. Abhishek Tanwar: Formal analysis; Investigation. Anuradha Jadon: Formal analysis. Ankit Biswas: Formal analysis; Investigation Tushar K Maiti: Resources; Investigation; Supervision. Rajender K Motiani: Conceptualization; Supervision; Funding acquisition; Writing—original draft; Project administration; Writing-review and editing.

## Methods-

### Cell Culture

B16-F10 mouse melanoma cells were procured from the ATCC and cultured in Dulbecco’s Modified Eagle’s Medium-high glucose (DMEM-HG; Sigma-Aldrich, D5648) supplemented with 10% fetal bovine serum (FBS; Gibco, 10270106) and 1X Antibiotic Antimycotic (Thermo Fisher Scientific, 15240062). Cells were maintained at 60-80% confluency in a humidified incubator at 5% CO_2_ levels and 37° C. Lightly pigmented (LP) neonatal primary human melanocytes were obtained from Invitrogen, Waltham, Massachusetts, USA. Cells were grown in Medium-254 (M254; Gibco, M254CF) supplemented with human melanocyte growth supplement-2 (HMGS-2; Gibco, S0165) and 1X Antibiotic Antimycotic (Thermo Fisher Scientific,15240062) under the same incubating conditions. Manufacturer’s instructions were followed for maintenance and sub-culturing of these cells. Cells from passages 2-6 were used for experimentation.

### B16 pigmentation models

For LD (low density) pigmentation experiments, non-pigmented B16 cells were seeded at a density of 100 cells/cm^2^ in DMEM-high glucose media supplemented with 10% FBS and allowed to pigment autonomously for 6 days as described earlier [26].

For HD (high density) pigmentation experiments, B16 cells were seeded followed by pigmentation induction using inducers – 1μM melanocyte-stimulating hormone (αMSH; Sigma Aldrich, M4135), 20μM Forskolin (Sigma Aldrich, F3917-10MG) and 60μM IBMX (3-Isobutyl-1-methylxanthine, Merck, I5879-100MG) for 48h-72h.

### Transfections

Transfections in B16 LD pigmentation model was performed on Day 3 after cell seeding (Day 0) in T75 flasks using SmartPool siRNAs from Dharmacon at 100nM working concentration (see Table). Transfection mix was made in 1:3 V:V ratio of siRNA and Dharmafect transfection reagent, respectively, in Opti-MEM (Gibco, 31985070) media which was then incubated over the cells for 4-6h, followed by its removal and subsequent addition of Day3 media back to the flasks.

For transfections in B16 HD pigmentation models, cells were seeded at a density of 2.0 × 10^4^ per well in 6-well plate. The following day, siRNA transfection mix was made using 50nM siRNA from Dharmacon and Eurogentec in 1:3 V:V ratio with TurboFect Transfection Reagent (Thermo Fisher Scientific, R0531) in serum free DMEM-HG, incubated for 10 min and added over the cells in complete media along with pigmentation inducing agent. Cells were harvested 72hpt and used for downstream experimentation.

For siRNA transfections in primary human melanocytes, Nucleofection kit from Lonza was used (VPD-1003, U-024 program). siRNAs from Dharmacon were used at a working concentration of 100nM. Cells were harvested 72hpt for phenotype observation.

Details of siRNAs used-

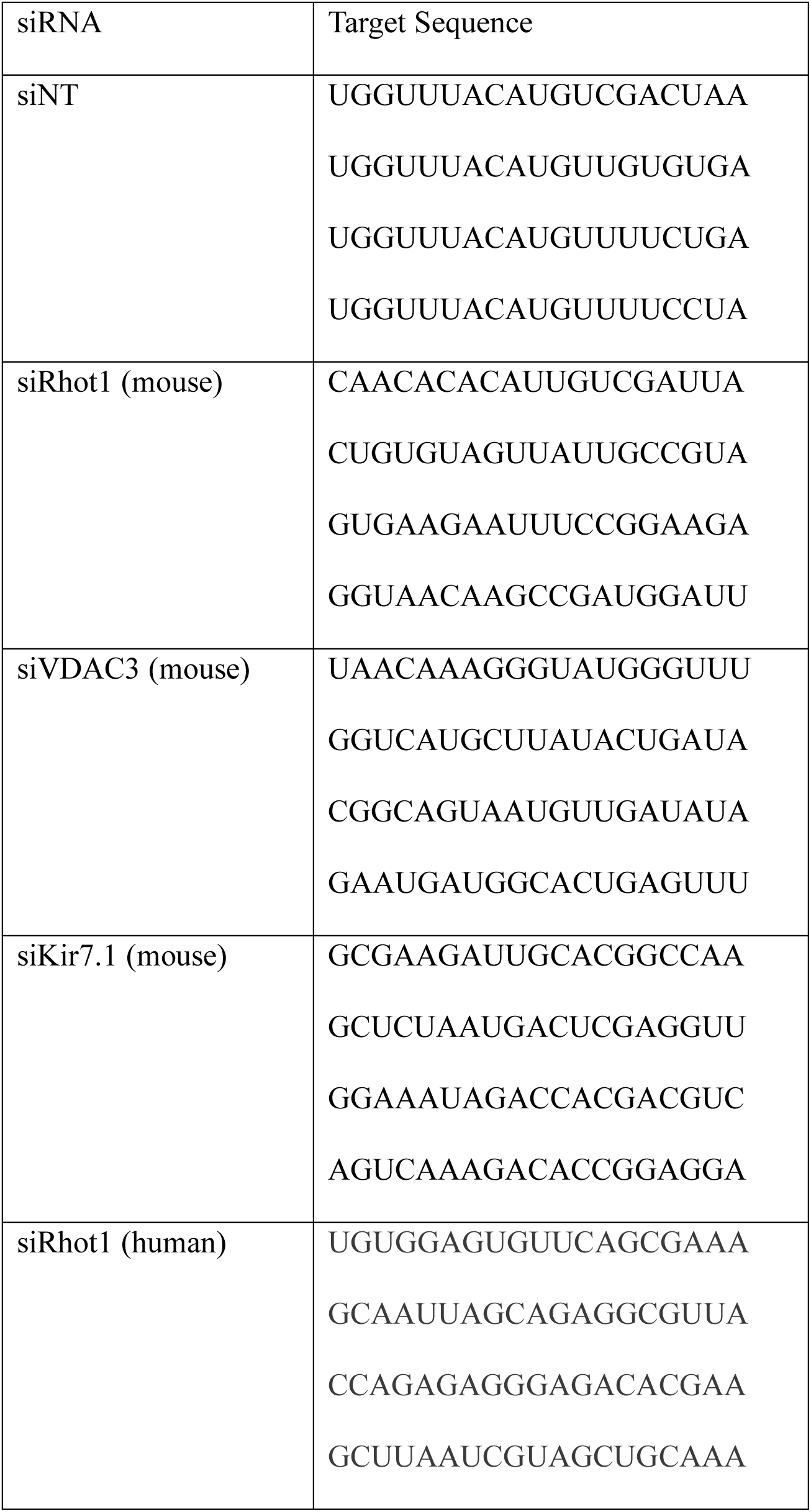
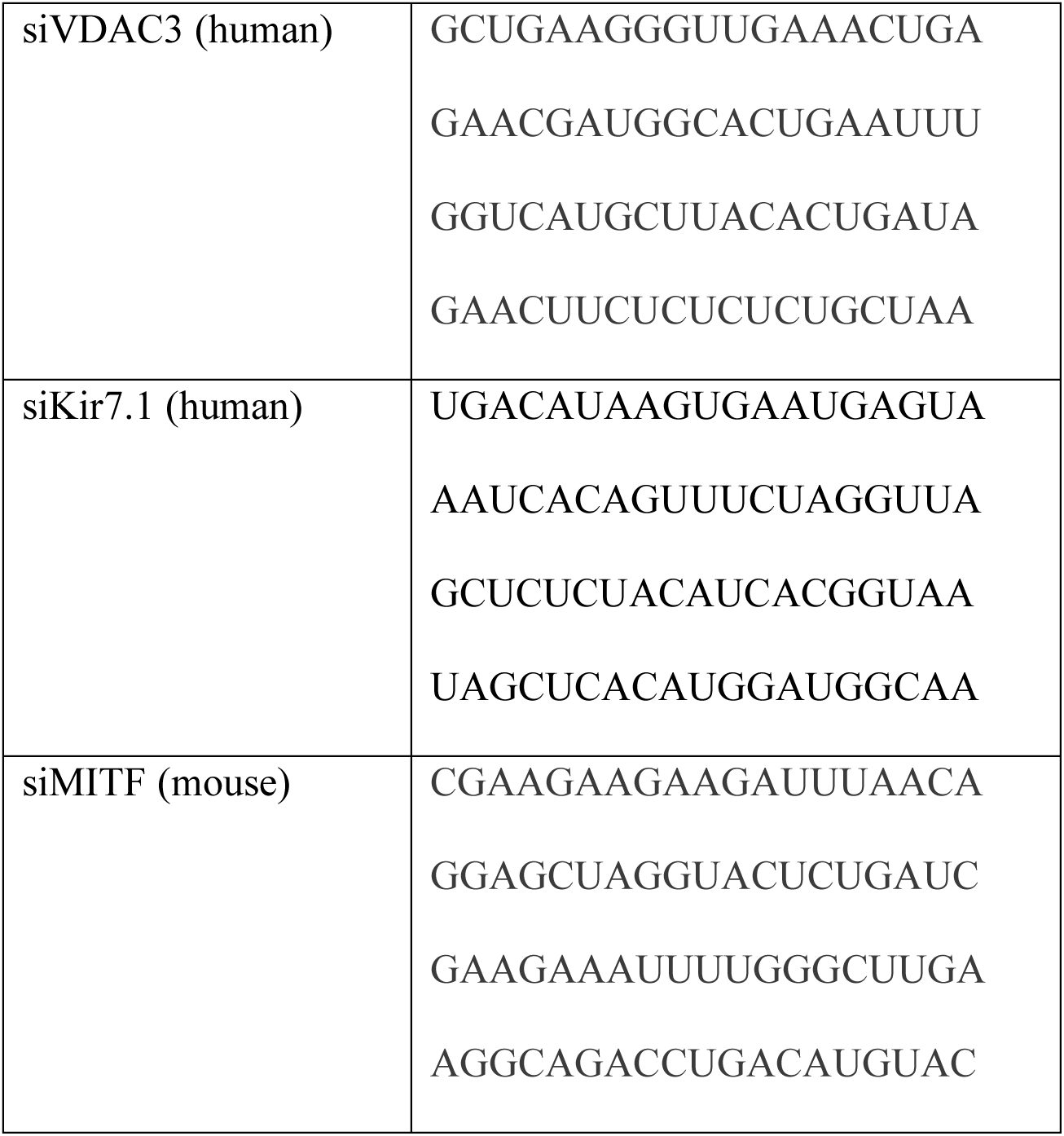

### Protein Over-expression in B16 cells

B16 cells were plated at a density of 5.0 × 10^4^ cells per well followed by plasmid transfection (0.5-2μg plasmid) using Turbofect transfection reagent (Thermo Fisher Scientific, R0531) in 1:2 V:V ratio. Pigmentation was induced using 1μM αMSH (Sigma Aldrich, M4135). Cells were harvested 48hpt. pRK5-myc-Miro1 was a gift from Pontus Aspenström (Addgene plasmid # 47888). pC1GFP-Kir7.1 and c1GFP-W53XKir7.1 were a gift from Bikash Pattnaik (Addgene plasmid # 79118 and # 107183 respectively).

| **Plasmid** | **Concentration Used** | **Source** | **Catalog Number** |
|---|---|---|---|
| pRK5-myc-Miro1 | 2μg | Addgene | #47888 |
| pC1GFP-Kir7.1 | 0.5μg | Addgene | #79118 |
| c1GFP-W53XKir7.1 | 0.5μg | Addgene | #107183 |

### qRT-PCR Analysis

Total cellular mRNA was extracted using the RNeasy Mini Kit (Qiagen, 74106). cDNA was then synthesized from the extracted RNA using High-Capacity cDNA Reverse Transcription Kit (Thermo Fisher Scientific, 4368814). Quantitative real time-PCR reactions were set up using SYBR green (Takara, RR420A) in Applied Biosystems Quant Studio 6 Flex or Bio-Rad CFX96 Touch Real-Time PCR Detection System. Relative mRNA levels were calculated using the ΔΔCt method with GAPDH as the normalization control unless otherwise stated. Gene specific primers are enlisted in Table -

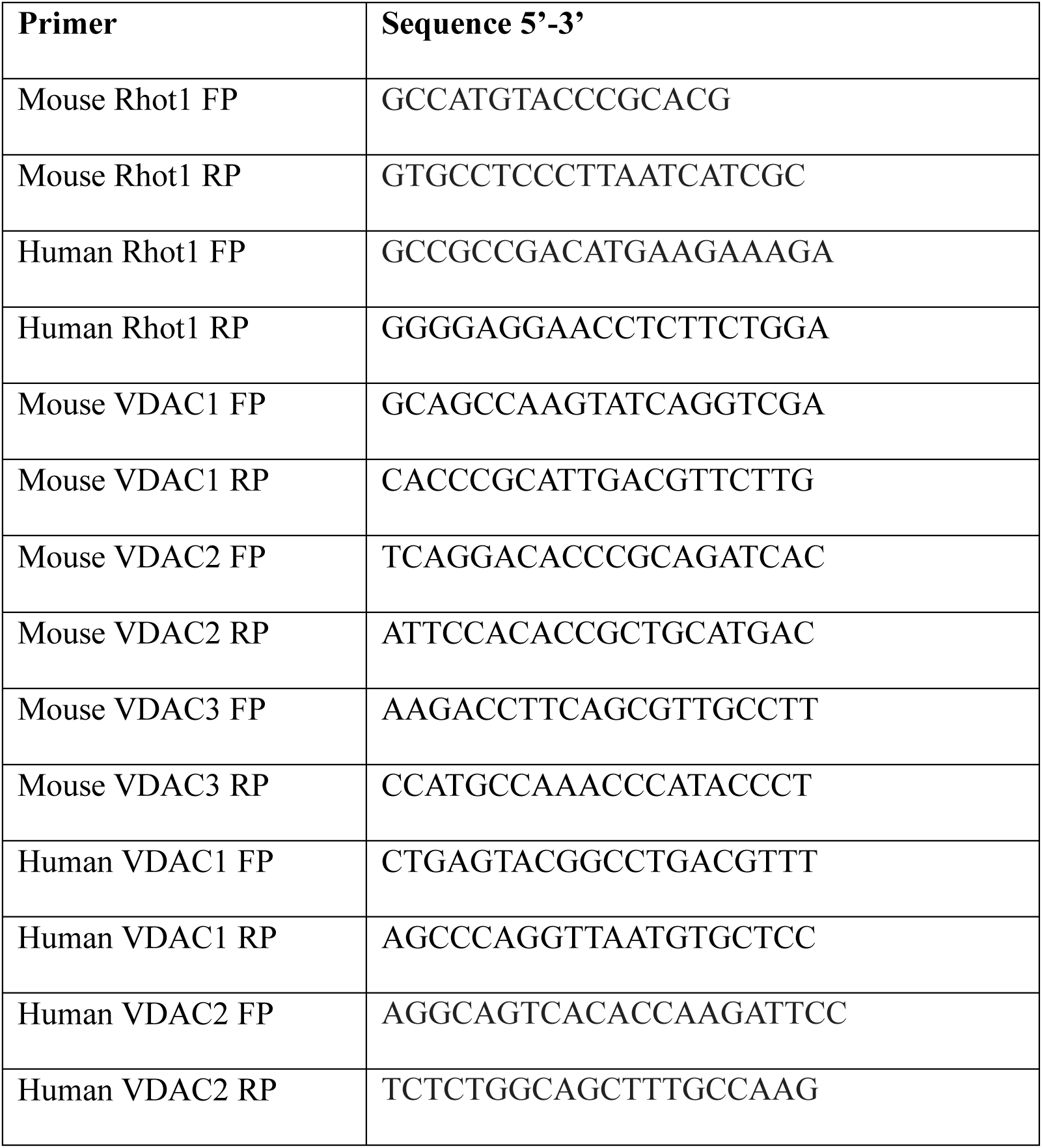
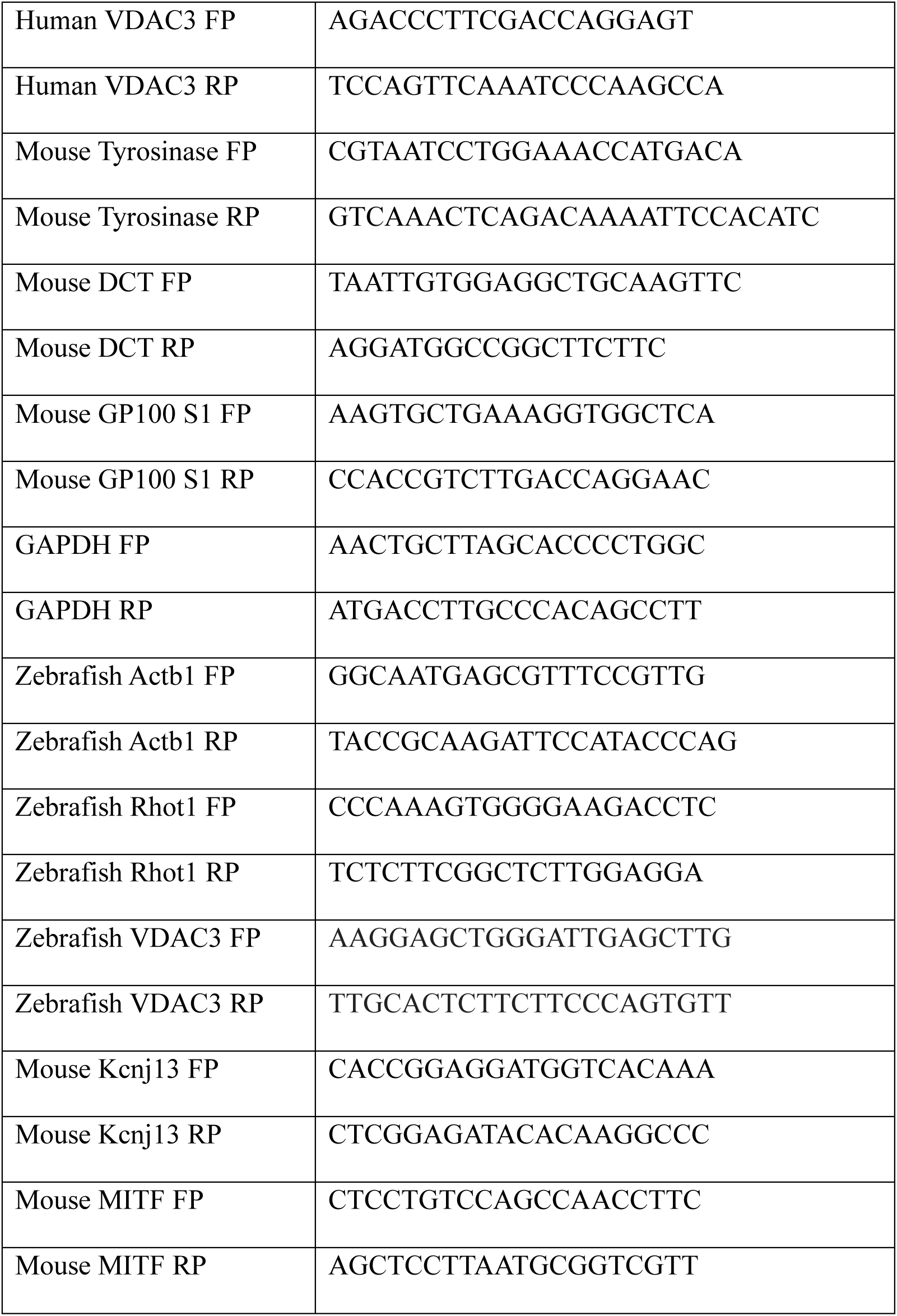

### Western Blotting

Cells were lysed using NP40 lysis buffer (Gibco, FNN0021) supplemented with Protease Inhibitor Cocktail (Roche, 4693159001). Protein concentration was quantified using the Pierce BCA Protein Assay Kit (Thermo Fisher Scientific, 23227) according to the manufacturer’s instructions. Typically, 20-50μg proteins were subjected to SDS-PAGE using 10-12% gel, followed by transfer on to a PVDF membrane. PVDF membranes were blocked in Tris-buffered saline containing 0.1% Tween 20 (TBST) and 5% skimmed milk for 2h at RT and incubated with primary antibodies diluted typically at 1:500 to 1:2,000 in 2% skimmed milk prepared in TBST and incubated overnight at 4 °C. Following day, after 3 washes, membrane was incubated in horseradish-peroxidase-conjugated anti-mouse or anti-rabbit IgG antibodies for 2h at RT [61]. Protein bands were visualized using Immobilon Forte Western HRP substrate (Millipore, WBLUF0500) and acquired using the ImageQuant LAS4000 imaging system (GE Healthcare). Densitometric analysis of protein bands was performed using FIJI software.

List of antibodies used-

| **Antibody** | **Manufacturer** | **Catalog Number** |
|---|---|---|
| VDAC3 polyclonal | ProteinTech | 55260-1-AP |
| DCT | Abcam | ab74073 |
| GP100 | Abcam | ab137078 |
| Tyrosinase | Invitrogen | 35-6000 |
| β-Tubulin | Abcam | ab21058 |
| Kir7.1 Antibody (C-12) | Santa Cruz | sc-398810 |
| GAPDH | Cell Signaling Technology | CST2118 |
| β-Actin | Cell Signaling Technology | CST4967 |
| Calnexin | Abcam | ab22595 |
| Phospho-PKA Substrate (RRXS*/T*) | Cell Signaling Technology | CST9624 |

### Mitochondrial Calcium Imaging

For mitochondrial Ca^2+^ imaging, B16 cells (0.6 ×10^5^ cells/well) were seeded on the confocal dishes (SPL Life Sciences, Korea), followed by siRNA transfections on the next day. 24h before imaging time point, pCMV-CEPIA2mt plasmid (Addgene #58218; 1.5 µg; excitation/emission: 488 nm/500-550 nm) was transfected using Lipofectamine 2000 (Invitrogen, 11668–019) in 1:2 (plasmid:transfection reagent) ratio. On the imaging day, cells were washed 3 times and bathed in HEPES-buffered saline solution (140 mM NaCl, 1.13 mM MgCl_2_, 4.7 mM KCl, 2 mM CaCl_2_, 10 mM D-glucose, and 10 mM HEPES; pH 7.4) for 5 min before Ca^2+^measurements. Imaging was performed on Nikon Eclipse Ti2 microscope using 60X oil objective. During live cell imaging, cells were stimulated by adding 50µm Histamine in the bath solution to trigger IP3R mediated Ca^2+^ efflux from the ER. At least 3 independent experiments were performed and ‘n’ denotes the number of cells. Data was plotted using GraphPad Prism software. pCMV-CEPIA2mt was a gift from Masamitsu Lino (Addgene #58218).

### Zebrafish husbandry

Zebrafish used in this study were housed in the Laboratory of Calciomics and Systemic Pathophysiology at the Regional Centre for Biotechnology, India, in accordance with institutional ethical guidelines and with approval from the Institutional Animal Ethics Committee of Regional Centre for Biotechnology, India (Protocol No: RCB/IAEC/2021/086) ensuring minimal animal suffering.

### Morpholino design and microinjections

Translation blocking antisense morpholino (MO) oligonucleotides designed against zebrafish Rhot1 and VDAC3 genes were procured from Gene Tools, USA. Morpholinos were reconstituted using nuclease-free water to obtain 1mM stock. Zebrafish embryos collected were transferred to 200μM phenylthiourea (PTU) containing embryo water at 24hpf for 24h to attenuate pigment synthesis. Next day, Morpholinos were injected at the hindbrain ventricle region of the zebrafish larvae and the larvae were transferred to normal embryo water to allow progression of pigmentation. 50μM Rhot1 MO and 100μM VDAC3 MO dosage of injection were used for injections along with same concentrations of Scrambled MO injections used as negative control. Pigmentation phenotype of the embryos was recorded at 24h post injection using Nikon microscope. The data was quantitated using ImajeJ software.

Sequences of the morpholinos used-

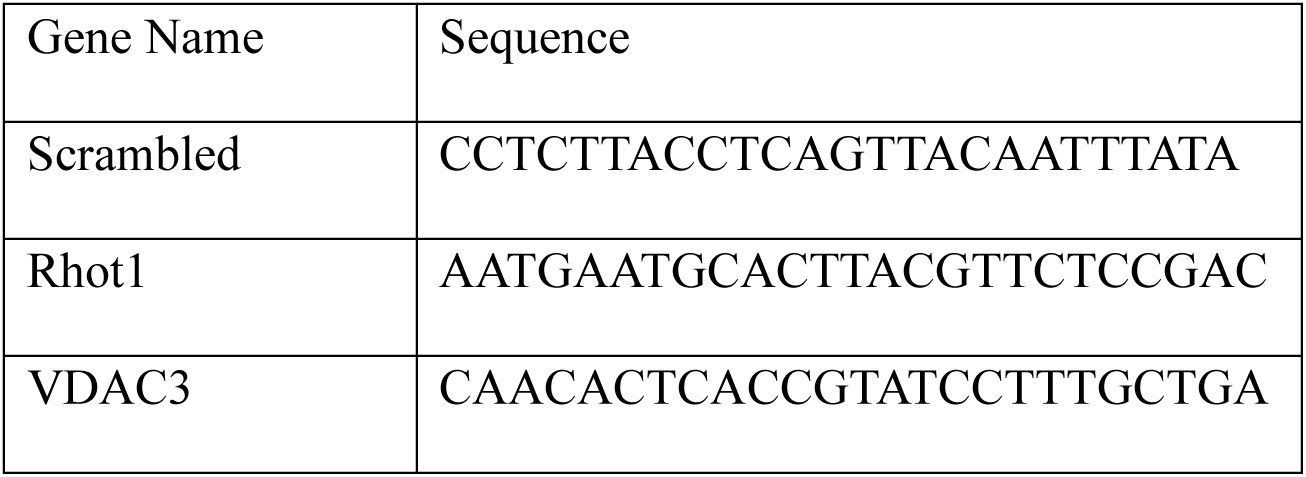

### Proteomics sample preparation, data acquisition and analysis

The transfected B16 cells are confirmed for the knock-down of Rhot1 and VDAC3, followed by their collection in RIPA (R0278, Sigma) buffer added with 1X halt protease phosphatase inhibitor (78441, Thermo Scientific). The harvested cells were lysed by probe sonication and then centrifuged to isolate the supernatant containing the total extracted proteins. The extracted proteins were precipitated using chilled acetone and then redissolved in 8M urea. The urea concentration is diluted till 2M using 100 mM ammonium bicarbonate (ABC) and finally the total protein concentration is measured using BCA protein assay kit (23225, Pierce, Thermo Scientific). For tryptic digestion, 50 ug of protein was reduced with 10 mM DL-Dithiothreitol (DTT) (D5545, Sigma) at 56 °C for 45 mins, and then alkylated with 20 mM Iodoacetamide (IAA) (I1149, Sigma) for 1 hour at room temperature. These reduced and alkylated samples were digested with a 1:20 (E:S) ratio of MS-grade trypsin (90057, Pierce, Thermo Scientific) at 37 °C. After, overnight incubation the tryptic peptides were passed through Oasis HLB C18 cartridges (WAT094225, Waters) using an Extraction Manifold (WAT200607, Waters, USA) to remove the salts. The samples were vacuum dried and stored at -80 °C until ready for MS acquisition.

The samples were reconstituted in solvent A (2% [vol/vol] acetonitrile, 0.1% [vol/vol] formic acid in water) and then loaded onto Luna 5 µm C18 Micro trap column (20 mm X 0.3 mm, 100 Å, Phenomenex) which is connected with nanoEase M/Z HSS T3 analytical column (150 mm X 300 µm, 1.8 µm, 100 Å, Waters) for sample separation. The binary pump module on waters microscale LC system (ACQUITY UPLC M-Class System) was used for resolving the injected (500 ng) peptides over a linear gradient of solvent B (100% (v/v) acetonitrile with 0.1% (v/v) formic acid) at 5 µL/min flowrate. The data were acquired in ZenoSWATH mode using ZenoTOF 7600 (Sciex) mass spectrometer utilizing 65 variable sized overlapping windows.

Post data acquisition, the .wiff files were used for de-novo spectral library generation in Spectronaut pulsar 19 (Biognosys) against the UniProtKB human protein database with isoforms (42,502 entries, December 2024) and default settings. Next, this spectral library was utilized for protein group identification and differential abundance calculation with standard parameters. Briefly, trypsin is used as specific enzyme along with selected fixed modification (cysteine carbamidomethylation, +57.0215 Da) and variable modifications (methionine oxidation, +15.9949 Da; N-terminal acetylation, +42.0106 Da). A 1% false discovery rate cutoff was used at precursor, peptide and protein level for robust identification of protein groups. The global TIC (total intensity count) normalized data were corrected at MS2 level for any interference to isolate the extracted ion chromatogram (XIC) of the peptides which were further grouped to quantify the protein abundance in each samples. The PGquant values along with fold change and significance were extracted from the software for the downstream analysis in R (R 4.3.1). The differential proteins were used for gene set enrichment analysis (GSEA) using ClusterProfiler (v4.12.6) against ReactomePA (1.48.0) database.

### KCl Supplementation Assay

B16 cells were seeded 24 h before treatment at a density of 3.0 × 10^4^ cells/well in DMEM supplemented with 10% FBS in 6-well plates. In B16 cells, seeded at HD (High Density), pigmentation was induced by adding 1µM αMSH (Sigma-Aldrich, M4135) for 48h. Cell culture media was then supplemented with 0,10,25mM KCl for 48h followed by termination and cell pellet preparation. Mean pixel intensity of pellets was calculated using FIJI software.

### Valinomycin Mediated Pigmentation Assay

B16 cells were seeded 24h before treatment at a density of 5.0 × 10^4^ cells/well in DMEM supplemented with 10% FBS in 6-well plates. Pigmentation was induced by adding 1µM αMSH (Sigma-Aldrich, M4135) for 48h. Cells were then treated with 2nM Valinomycin (Sigma-Aldrich, V0627-10MG) with DMSO as control for each concentration for 48h followed by termination and cell pellet preparation. Mean pixel intensity of pellets was quantitated using FIJI software.

### VU590 Mediated Pigmentation Assays

B16 cells were seeded 24 h before treatment at a density of 5.0 × 10^4^ cells/well in DMEM supplemented with 10% FBS in 6-well plates. Next day, pigmentation was stimulated in cells using 1μM melanocyte-stimulating hormone (αMSH; Sigma Aldrich, M4135), 20μM Forskolin (Sigma Aldrich, F3917-10MG) or 60μM IBMX (3-Isobutyl-1-methylxanthine, Merck, I5879-100MG). The following day, cells were treated with 8,10 and 15μM VU590 (MedChemExpress, HY-108595). Experiment was terminated 24h post drug treatment and cell pellets were prepared.

For combinatorial VU590 and KCl treatment experiments, B16 cells were seeded 24 h before treatment at a density of 2.5 × 10^4^ cells/well in DMEM supplemented with 10% FBS in 6-well plates. In B16 cells, seeded at HD, pigmentation was induced by adding 20uM forskolin (Sigma-Aldrich, F3917) for 48h. Cells were then treated with 8uM VU590 with DMSO as control. After 6 h, 0,10,25 mM KCl was added to media for 42 hours followed by termination and cell pellets were made. Mean pixel intensity of pellets was quantified using FIJI software.

### H-89 Mediated Pigmentation Assays

B16 cells were seeded at a density of 2.0 × 10^4^ cells/well in DMEM supplemented with 10% FBS in 6-well plates. Next day, cells were transfected with siRNAs using the mentioned protocol. 24 hours after transfection, cells were pre-treated with 1uM H-89 and control cells were pre-treated with the same volume of DMSO for 1 hour incubated at 37°C and 5% CO_2_. This was followed by replacement with media containing 1uM H-89 along with 1μM melanocyte-stimulating hormone (αMSH; Sigma Aldrich, M4135), or DMSO with 1μM αMSH in control conditions. The experiment was terminated 48 hours post drug treatment and cell pellets were made for further analyses.

### Transmission Electron Microscopy (TEM)

Cells were harvested and resuspended in phosphate buffer (PB), pelleted by centrifugation, and fixed in TEM fixative for 2h at 4 °C. Following fixation, samples were centrifuged to remove the fixative and resuspended in PB. Specimens were processed for transmission electron microscopy, and ultrathin sections (60–90 nm) were prepared. Sections were double-stained with uranyl acetate and alkaline lead citrate. TEM images were acquired using a JEM-1400 Flash transmission electron microscope equipped with a tungsten filament electron source and a high-sensitivity sCMOS camera, operated at 80 kV and a magnification of 30,000X.

### Intracellular Potassium Imaging

For comparative visualization of intracellular K^+^ levels, cells were stained using 5µM ION Potassium Green-2 AM (Abcam, ab142806) for 30 min at 37°C and 5% CO_2_ levels. After washing twice with PBS, cells were subjected to confocal microscopy (excitation/emission 526/546nm) on Zeiss LSM 880 confocal microscope using 63X oil objective. The data was analysed using FIJI software.

### Cell Membrane Potential Measurement

Cell membrane potential was measured with DiBAC4(3) (Bis-(1,3-Dibutylbarbituric Acid Trimethine Oxonol, Thermo Fischer Scientific, B438). B16 cells were loaded with 1µm dye in 1X HBSS (HEPES-buffered saline solution: 140 mM NaCl, 1.13 mM MgCl2, 4.7 mM KCl, 10 mM D-glucose, and 10 mM HEPES; pH 7.4) for 30 min at 37°C and 5% CO_2_ levels followed by confocal imaging (excitation/emission: 493/516nm) on Zeiss LSM 880 confocal microscope using 63X oil objective. The data was analysed using FIJI software.

## Database Analysis

The publicly accessible microarray dataset (accession IDs: GSE53146, GSE65127, GSE72140) were analysed using the GEO2R tool available at the microarray data repository Gene Expression Omnibus (https://www.ncbi.nlm.nih.gov/geo/geo2r) of NCBI. Whole-skin gene expression levels of the *Kcnj13* gene were compared between the control and test groups. Normalized expression values after log transformation are visualised using the Graphpad Prism.

## Statistical analyses

Statistical analyses were performed using ANOVA, Student’s paired or unpaired t-test, one sample t-test as appropriate using GraphPad Prism. P values <0.05, <0.01, <0.001 and <0.0001 are identified with *, **, *** and ****, respectively. "ns” indicates P ≥ 0.05.

---

**Supplementary Figure 1 supporting Main Figure 1.**
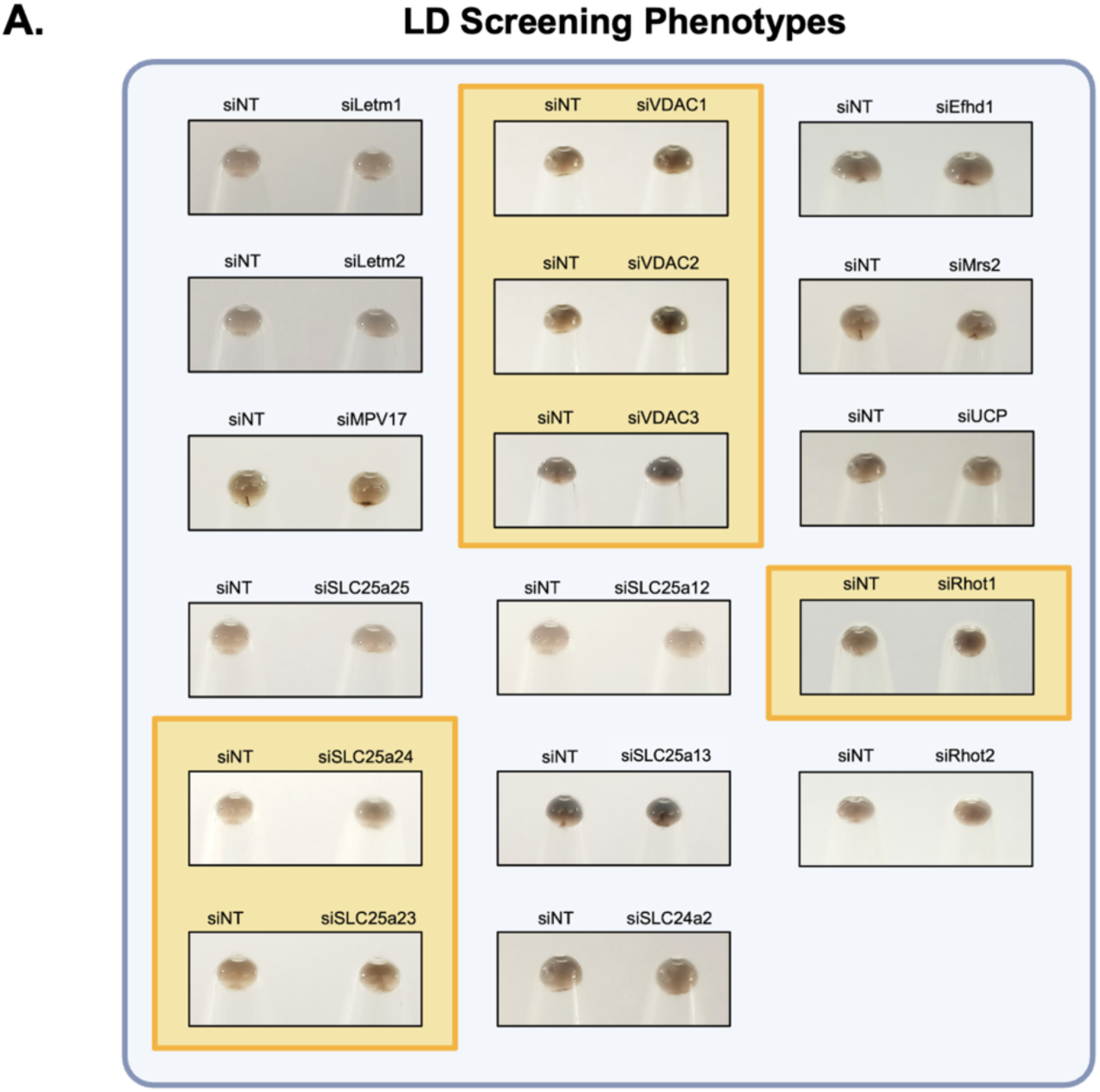
Unbiased siRNA screens targeting mitochondrial Ca^2+^ handling machinery identify novel modulators of pigmentation. A. Pigmentation phenotypes upon knockdown of target candidate proteins in LD day 6 B16 cells as compared to respective siNT controls.

**Supplementary Figure 2 supporting Main Figure 2.**
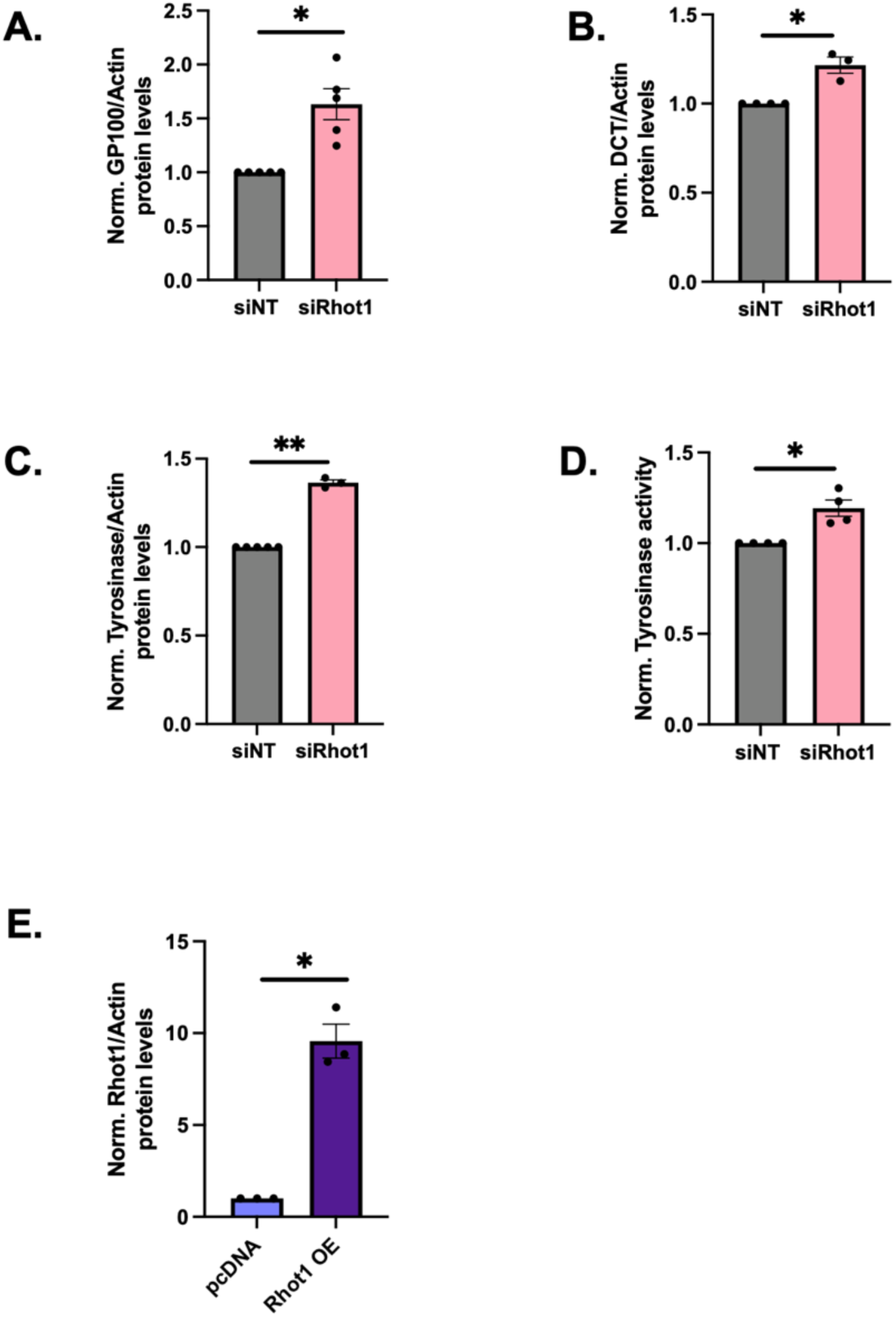
Rhot1 is a negative modulator of melanogenesis. A. Densitometric quantitation showing increase in GP100 levels in siRhot1 LD day 6 cells as compared to siNT control (N=5). B. Densitometric quantitation showing increase in DCT levels in siRhot1 LD day 6 cells as compared to siNT control (N=3). C. Densitometric quantitation showing increase in tyrosinase levels in siRhot1 LD day 6 cells as compared to siNT control (N=3). D. DOPA assay showing increased tyrosinase activity in siRhot1 LD day 6 cells as compared to siNT control (N=4). E. Densitometric quantitation of Rhot1 protein levels in Rhot1 OE condition compared to the control condition (N=3). Data presented are mean ± SEM. For statistical analysis, one sample t and Wilcoxon test was performed for panels A-E using GraphPad Prism software. Here, * p<0.05; ** p<0.01; *** p<0.001; and **** p<0.0001.

**Supplementary Figure 3 supporting Main Figure 3.**
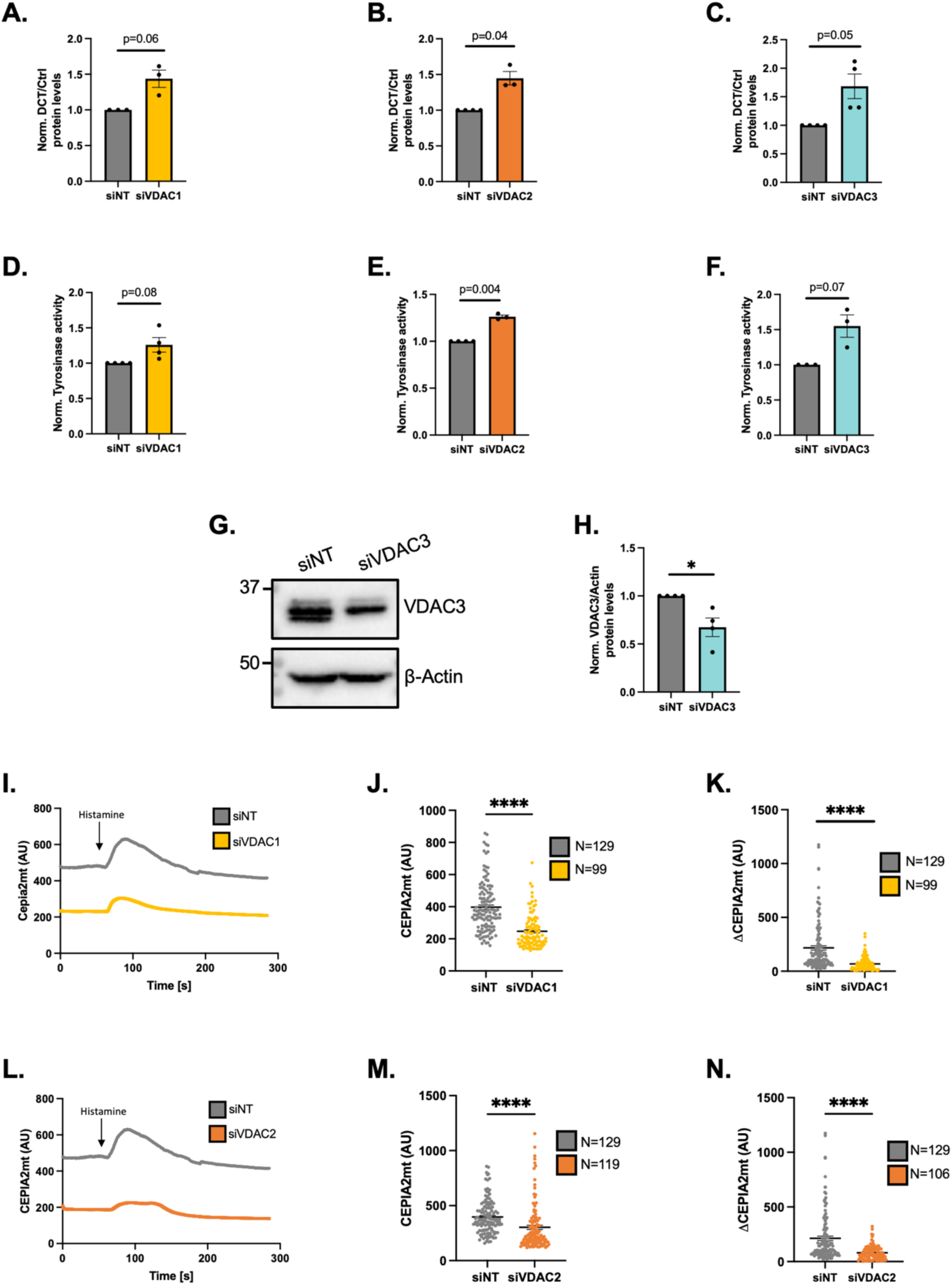
VDAC3 is a negative modulator of melanogenesis. A. Densitometric quantitation showing increase in DCT levels in siVDAC1 LD day 6 cells as compared to siNT control (N=3). B. Densitometric quantitation showing increase in DCT levels in siVDAC2 LD day 6 cells as compared to siNT control (N=3). C. Densitometric quantitation showing increase in DCT levels in siVDAC3 LD day 6 cells as compared to siNT control (N=4). D. Densitometric quantitation showing tyrosinase activity in siNT control and siVDAC1 LD day 6 cells (N=4). E. Densitometric quantitation showing tyrosinase activity in siNT control and siVDAC2 LD day 6 cells (N=3). F. Densitometric quantitation showing tyrosinase activity in siNT control and siVDAC3 LD day 6 cells (N=3). G. Representative western blot validating VDAC3 knock-down in siVDAC3 condition compared to siNT control (N=3). H. Densitometric analysis confirming VDAC3 silencing in siVDAC3 B16 cells compared to siNT control (N=3). I. Representative mitochondrial Ca^2+^ imaging traces of CEPIA2mt in siNT control and siVDAC1 B16 cells stimulated with 50µM histamine. J. Quantitation of resting mitochondrial Ca^2+^ with CEPIA2mt in siNT control and siVDAC1 B16 cells where “N” denotes the number of ROIs. K. Quantitation of mitochondrial Ca^2+^ uptake by calculating increase in CEPIA2mt signal (ΔCEPIA2mt) in siNT control and siVDAC1 B16 cells stimulated with 50µM histamine where “N” denotes the number of ROIs. L. Representative mitochondrial Ca^2+^ imaging traces of CEPIA2mt in siNT control and siVDAC2 B16 cells stimulated with 50µM histamine. M. Quantitation of resting mitochondrial Ca^2+^ with CEPIA2mt in siNT control and siVDAC2 B16 cells where “N” denotes the number of ROIs. N. Quantitation of mitochondrial Ca^2+^ uptake by calculating increase in CEPIA2mt signal (ΔCEPIA2mt) in siNT control and siVDAC2 B16 cells stimulated with 50µM histamine where “N” denotes the number of ROIs. Data presented are mean ± SEM. For statistical analysis, one sample t and Wilcoxon test was performed for panels A, B, C, D, E, F and H, while unpaired Student’s t test was performed for panels J, K, M and N, using GraphPad Prism software. Here, * p<0.05; ** p<0.01; *** p<0.001; and **** p<0.0001.

**Supplementary Figure 4 supporting Main Figure 5.**
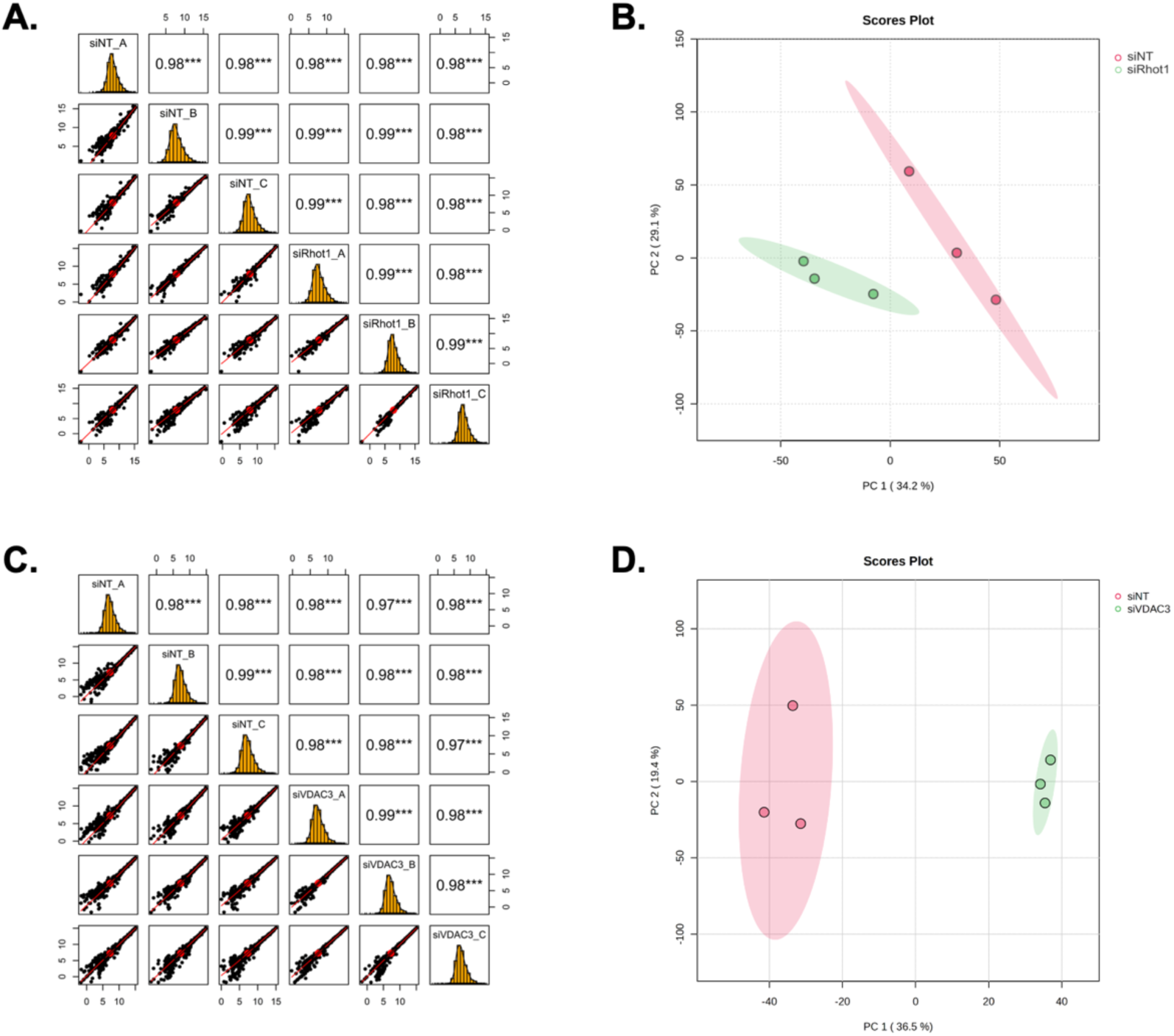
Proteomics analysis identifies Kir7.1 functioning downstream of Rhot1 and VDAC3 to modulate melanogenesis. A. Quality control (QC) metrics of siNT vs siRhot1 HD-MSH B16 cells dataset (N=3). B. Principal component analysis (PCA) of siNT vs siRhot1 HD-MSH B16 cells dataset (N=3). C. Quality control (QC) metrics of siNT vs siVDAC3 HD-MSH B16 cells dataset (N=3). D. Principal component analysis (PCA) of siNT vs siVDAC3 HD-MSH B16 cells dataset (N=3).

**Supplementary Figure 5 supporting Main Figure 6.**
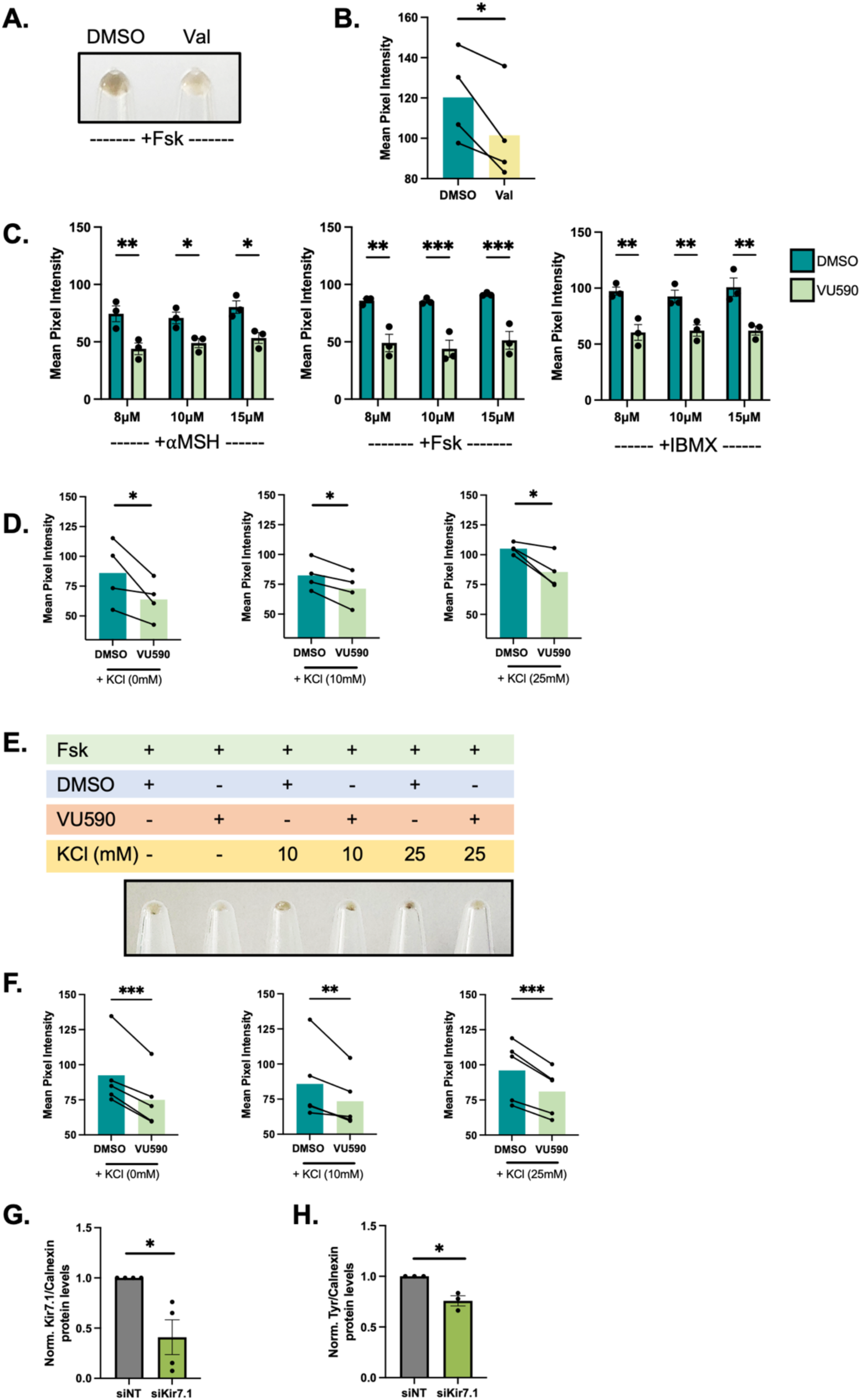
Kir7.1 positively modulates pigmentation. A. Representative pellet picture showing reduced pigmentation phenotype upon treating B16 cells with Valinomycin as compared to DMSO vehicle control. B. Mean pixel intensity-based pigmentation phenotype quantification upon treating B16 cells with Valinomycin. (N=4). C. Mean pixel intensity-based pigmentation phenotype quantification upon VU590 treatment in B16 cells stimulated with different melanogenic stimulators-αMSH, forskolin and IBMX (N=3). D. Mean pixel intensity-based pigmentation phenotype quantification upon combinatorial treatment of increasing KCl concentrations and VU590 in B16 cells stimulated with αMSH (N=4). E. Representative pellet picture showing pigmentation phenotype upon combinatorial treatment of increasing KCl concentrations and VU590 on B16 cells stimulated with forskolin. F. Mean pixel intensity-based pigmentation phenotype quantification upon combinatorial treatment of increasing KCl concentrations and VU590 in B16 cells stimulated with forskolin (N=5). G. Densitometric quantitation confirming Kir7.1 knockdown in siKir7.1 in primary human melanocytes as compared to siNT control (N=4). H. Densitometric quantitation of tyrosinase expression upon Kir7.1 silencing in primary human melanocytes (N=3). Data presented are mean ± SEM. For statistical analysis, ratio paired t test was performed for panels B, D and F, two-way ANOVA + Šídák multiple-comparisons test was performed for panel C, while one sample t and Wilcoxon test was performed for panels G and H, using GraphPad Prism software. Here, * p<0.05; ** p<0.01; *** p<0.001; and **** p<0.0001.

**Supplementary Figure 6 supporting Main Figure 7.**
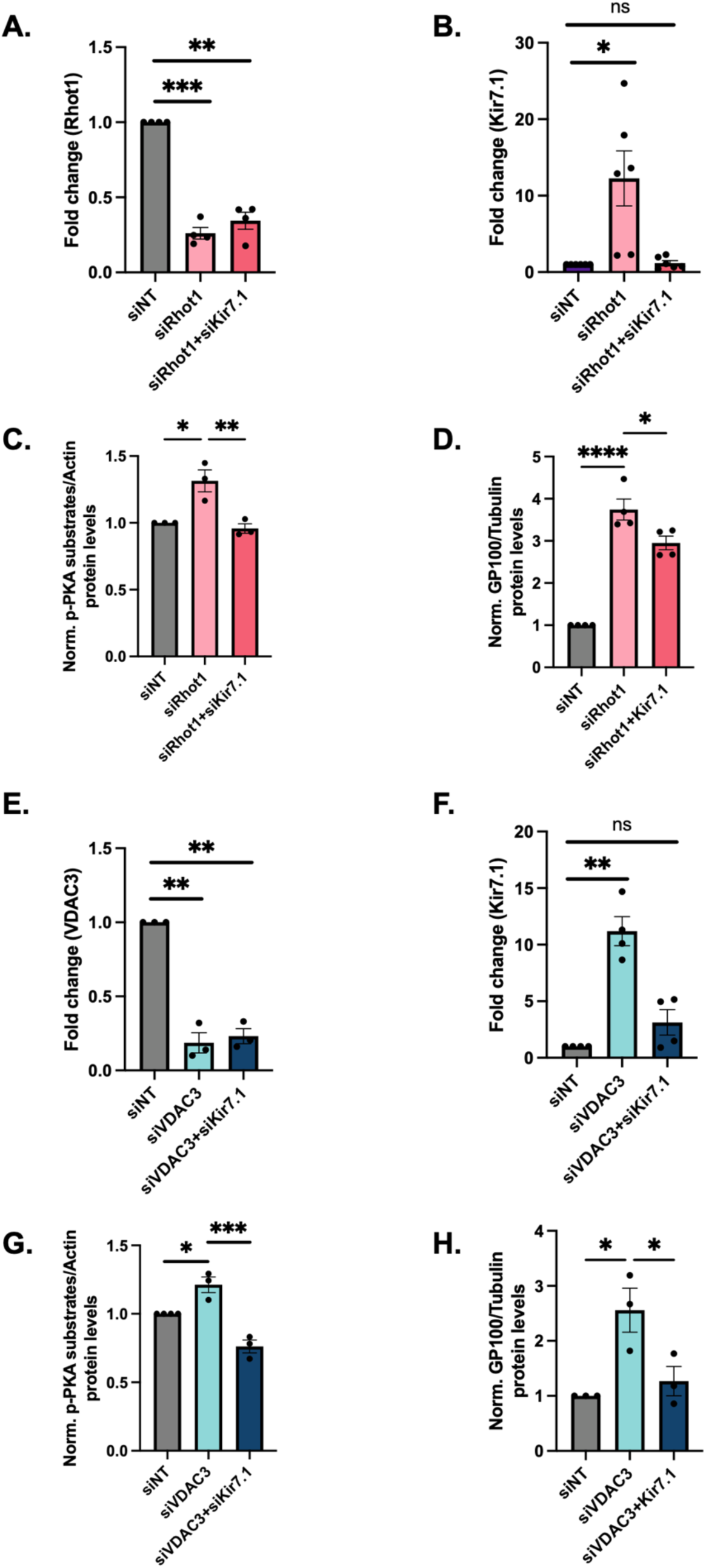
Kir7.1 functions downstream of Rhot1 and VDAC3 to modulate melanosome biogenesis and maturation. A. qRT-PCR analysis confirming siRNA based Rhot1 silencing in siRhot1 and siRhot1+siKir7.1 B16 cells (N=4). B. qRT-PCR analysis showing Kir7.1 mRNA levels in siRhot1 and siRhot1+siKir7.1 B16 cells (N=6). C. Densitometric quantitation showing phosphorylated PKA substrate levels in siRhot1 and siRhot1+siKir7.1 B16 cells as compared to siNT control (N=3). D. Densitometric quantitation showing GP100 levels in siRhot1 and siRhot1+siKir7.1 B16 cells as compared to siNT control (N=4). E. qRT-PCR analysis confirming siRNA based VDAC3 silencing in siVDAC3 and siVDAC3+siKir7.1 B16 cells (N=3). F. qRT-PCR analysis showing Kir7.1 mRNA levels in siVDAC3 and siVDAC3+siKir7.1 B16 cells (N=4). G. Densitometric quantitation showing phosphorylated PKA substrate levels in siVDAC3 and siVDAC3+siKir7.1 B16 cells as compared to siNT control (N=3). H. Densitometric quantitation showing GP100 levels in siVDAC3 and siVDAC3+siKir7.1 B16 cells as compared to siNT control (N=3). Data presented are mean ± SEM. For statistical analysis, one sample t and Wilcoxon test was performed for panels A, B, E, and F, while ANOVA was performed for panels C, D, G and H, using GraphPad Prism software. Here, * p<0.05; ** p<0.01; *** p<0.001; and **** p<0.0001.

**Supplementary Figure 7 supporting Main Figure 8.**
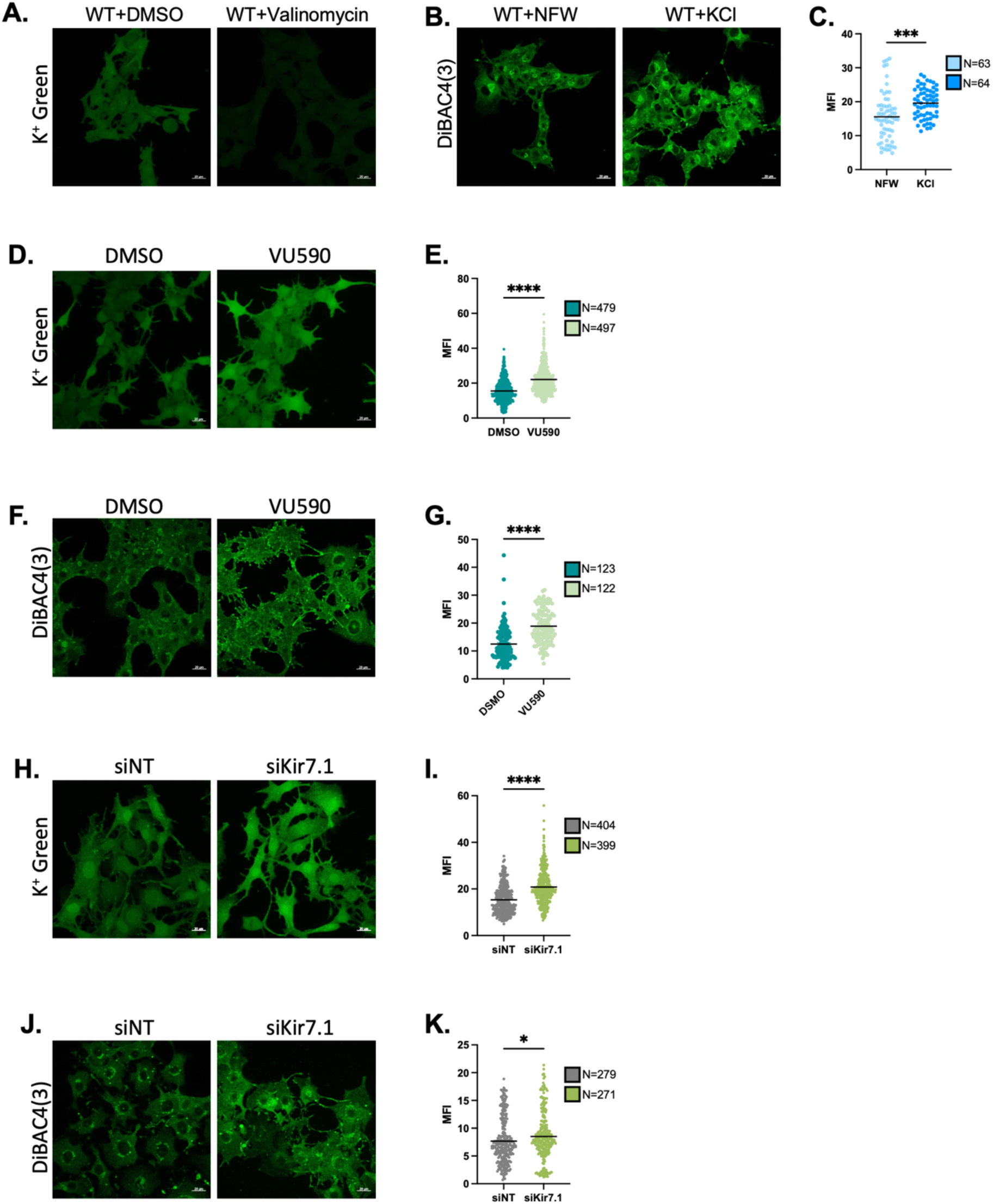
Kir7.1 affects cellular K^+^ homeostasis and membrane potential downstream of Rhot1 and VDAC3. A. Representative images for cellular K^+^ levels in DMSO and Valinomycin (K^+^ ionophore) treated B16 cells (Scale: 20μm). B. Representative images for cell membrane potential in NFW and KCl (depolarization control) treated B16 cells (Scale: 20μm). C. Quantification of mean fluorescence intensity indicating cell membrane potential in NFW and KCl (depolarization control) treated B16 cells, where ‘N’ denotes the number of cells imaged. D. Representative images for cellular K^+^ levels in DMSO and VU590 (Kir7.1 blocker) treated B16 cells (Scale: 20μm). E. Quantification of mean fluorescence intensity indicating cellular K^+^ levels in DMSO and VU590 treated B16 cells, where ‘N’ denotes the number of cells imaged. F. Representative images for cell membrane potential in NFW and VU590 treated B16 cells (Scale: 20μm). G. Quantification of mean fluorescence intensity indicating cell membrane potential in NFW and VU590 treated B16 cells, where ‘N’ denotes the number of cells imaged. H. Representative images for cellular K^+^ levels in siKir7.1 B16 cells compared to siNT control (Scale: 20μm). I. Quantification of mean fluorescence intensity indicating cellular K^+^ levels in siKir7.1 B16 cells compared to siNT control, where ‘N’ denotes the number of cells imaged. J. Representative images for cell membrane potential in siKir7.1 B16 cells compared to siNT control (Scale: 20μm). K. Quantification of mean fluorescence intensity indicating cell membrane potential in siKir7.1 B16 cells compared to siNT control, where ‘N’ denotes the number of cells imaged. Data presented are mean ± SEM. For statistical analysis, unpaired Student’s t-test was performed for panels C, E, G, I and K, using GraphPad Prism software. Here, * p<0.05; ** p<0.01; *** p<0.001; and **** p<0.0001.

## Notes

### Competing Interest Statement

The authors have declared no competing interest.

